# Comparing Functional Connectivity Matrices: A Geometry-Aware Approach applied to Participant Identification

**DOI:** 10.1101/687830

**Authors:** Manasij Venkatesh, Joseph Jaja, Luiz pessoa

## Abstract

Understanding the correlation structure associated with multiple brain measurements informs about potential “functional groupings” and network organization. The correlation structure can be conveniently captured in a matrix format that summarizes the relationships among a set of brain measurements involving two regions, for example. Such functional connectivity matrix is an important component of many types of investigation focusing on network-level properties of the brain, including clustering brain states, characterizing dynamic functional states, performing participant identification (so-called “fingerprinting”) understanding how tasks reconfigure brain networks, and inter-subject correlation analysis. In these investigations, some notion of proximity or similarity of functional connectivity matrices is employed, such as their Euclidean distance or Pearson correlation (by correlating the matrix entries). Here, we propose the use of a *geodesic distance* metric that reflects the underlying non-Euclidean geometry of functional correlation matrices. The approach is evaluated in the context of participant identification (fingerprinting): given a participant’s functional connectivity matrix based on resting-state or task data, how effectively can the participant be identified? Using geodesic distance, identification accuracy was over 95% on resting-state data, and exceeded the Pearson correlation approach by 20%. For whole-cortex regions, accuracy improved on a range of tasks by between 2% and as much as 20%. We also investigated identification using pairs of subnetworks (say, dorsal attention plus default mode), and particular combinations improved accuracy over whole-cortex participant identification by over 10%. The geodesic distance also outperformed Pearson correlation when the former employed a fourth of the data as the latter. Finally, we suggest that low-dimensional distance visualizations based on the geodesic approach help uncover the geometry of task functional connectivity in relation to that during resting-state. We propose that the use of the geodesic distance is an effective way to compare the correlation structure of the brain across a broad range of studies.

## 1. Introduction

Measurements of brain activity are acquired across multiple sensors or spatial locations, such as those obtained by electro/magneto-encephalography, electrophysiology recordings, calcium imaging, or functional magnetic resonance imaging (fMRI) data. Understanding the correlation structure associated with multiple brain measurements is a central goal in neuroscience, as it informs about potential “functional groupings” and network structure [29, 35]. The correlation structure can be conveniently captured in a matrix format that captures the relationships among a set of brain measurements. For example, in the case of fMRI, each entry of the matrix might contain an estimate of the *functional connectivity* (FC) between regions *i* and *j*, typically computed as the correlation between the time series data of the two regions in question.

In recent years, the FC matrix has become an important component of many types of investigation focusing on network-level properties of the brain, particularly in fMRI. For example, it has been used to cluster brain states [2], characterize dynamic functional states [19], perform participant identification [15], and understand how tasks reconfigure brain networks [33]. In these applications, some notion of proximity or similarity of FC matrices is employed (Fig. 1A). How should similarity be gauged? An intuitive approach is to “unroll” the FC matrix into a vector and compute the Pearson correlation between the matrices themselves. Thus if, say, two brain states captured by FC matrices are similar (for example, during two similar perceptual conditions), their matrices would be (relatively) highly correlated. Indeed, the correlation approach has yielded impressive results, such as successfully identifying a participant out of a large group of participants based on FC matrix similarity, a process dubbed fingerprinting ([15, 14, 3]). Related approaches include computing the Euclidean *(L*^2^) distance between the vectorized matrices [30], or using the so-called Manhattan (*L*^1^) distance [2].

**Fig. 1:**
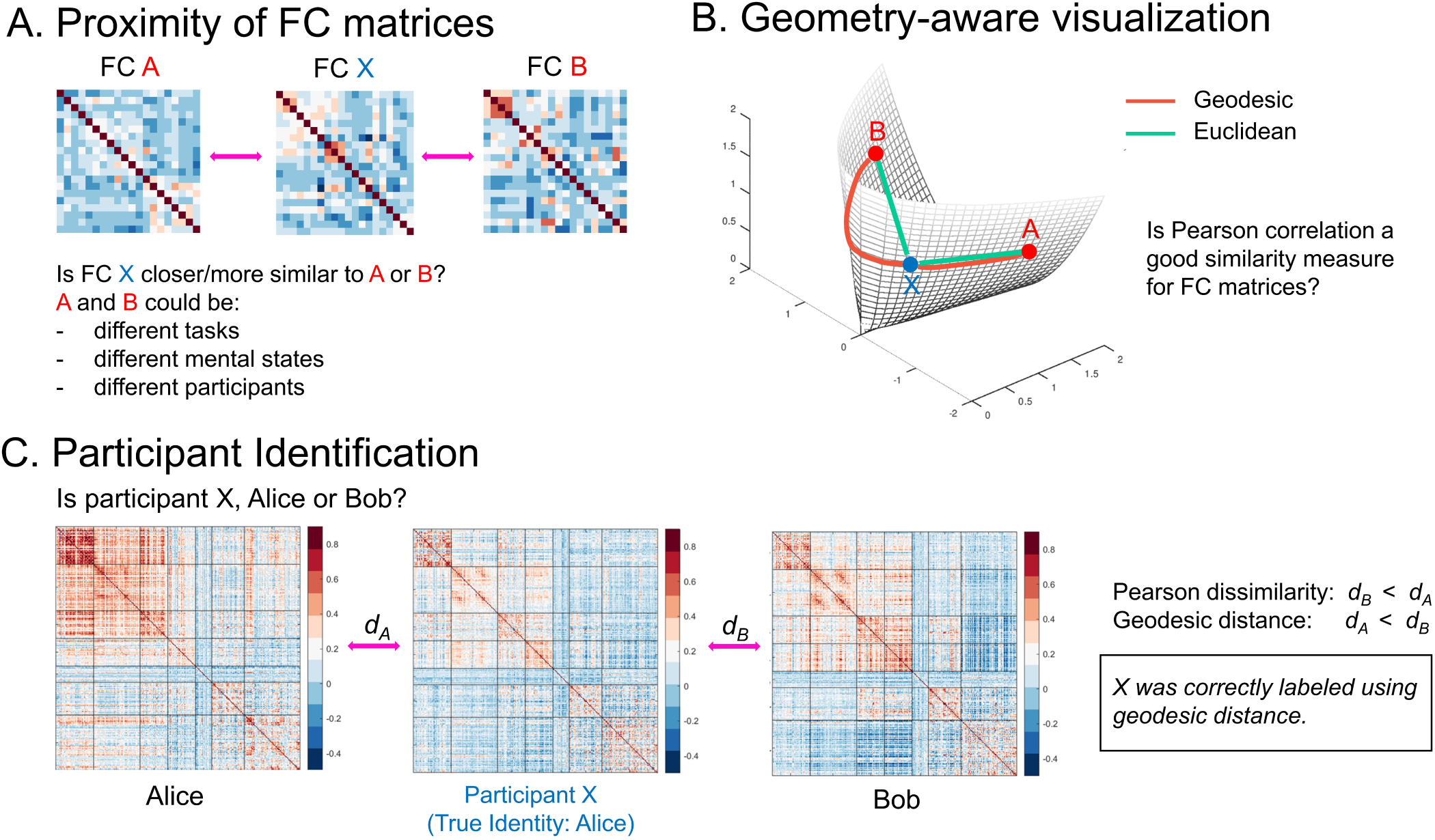
Functional connectivity matrices and their underlying geometry. (A) Similarity of functional connectivity (FC) matrices. Is the FC matrix *X* more similar to *A* or *B*? This question arises when the goal is to determine the task being being performed, the mental state, or the participant. (B) Illustration of geodesic distance (red) and Euclidean distance (green) on the so-called positive semidefinite cone. The geodesic and Euclidean distances between two points can differ substantially. (C) Is *X*, Alice or Bob? Equivalently, is the FC *X* more similar to that of Alice or Bob? Identification involves mapping an unknown participant’s data to one of the participants in the database (only two in this case). In this example, *X* is correctly labeled as Alice using geodesic distance, but incorrectly labeled as Bob using Pearson dissimilarity.

FC matrices computed by Pearson correlating time series data are objects that lie on a non-linear surface (technically known as a manifold) called the *positive semidefinite cone*: their geometry is non-Euclidean. Accordingly, distances between Pearson FC matrices must be measured along the surface of the cone (Fig. 1B). In addition, FC matrices are often high dimensional, and the proximity measure adopted is critical since noisy dimensions can contribute substantially to the measure [1].

In the present paper, we characterized the advantages of using a *geodesic* proximity measure between FC matrices. We apply the approach to the problem of participant identification: Given resting-state or task data, is it possible to determine a participant from her FC matrix [15]? We show that using the geodesic distance, a non-Euclidean distance metric that considers the manifold on which the data lies, improves participant identification compared to a similarity measure based on Pearson correlation (Fig. 1C). The improvement is shown to be non-trivial and consistent across resting-state and task conditions.

We also investigate how distances between high-dimensional FC matrices can be effectively visualized in low-dimensional spaces. Such visualization reflected identification accuracy based on the full-dimensional data, and thus retained important distance information. We suggest that visualization in lower dimensions aids in understanding the geometry of task FC structure in relation to resting-state FC.

## 2. Methods

### 2.1. Human Connectome Project Data

We utilized data from *N* = 100 unrelated participants from the Human Connectome Project (HCP) of the 1200-participant release [12]. Data from resting-state and seven tasks were employed: emotion processing (EM), gambling (GB), language (LG), motor (MT), relational processing (RL), social cognition (SO), and working memory (WM). Throughout the paper, we refer to resting-state plus the tasks as *conditions*. For a description of the tasks and scan parameters, see [5]. Data were collected with a repetition time (TR) of 720 ms.

During each run, stimuli were presented in separate blocks often interleaved with fixation blocks. Some task runs also contained cues. To retain only task-related segments of the run, extraneous segments were trimmed. To account for hemodynamic lag, the first four TRs of the block were not used, and the first four TRs following the end of the block were used [8]. Emotional processing, working memory, and motor tasks contained 3-second cues at block onset. Accordingly, to account for the cue response and the hemodynamic lag, data from 12 seconds after the cue onset to 3 seconds after the end of the block were used. Time course length for each condition before and after trimming is provided in Table 1. Note that trimming the fixation periods is important in characterizing participant identification from task data, because fixation periods behave much like “mini resting periods” that can potentially provide information regarding the participant. Analysis of data without trimming is included in supplemental material (Section S1).

**Table 1:** Number of frames per run (in samples) before and after trimming fixation periods.

| Condition | REST | EM | GB | LG | MT | RL | SO | WM |
| --- | --- | --- | --- | --- | --- | --- | --- | --- |
| Frames per full run | 1200 | 176 | 253 | 316 | 284 | 232 | 274 | 405 |
| Frames per trimmed run | 1200 | 141 | 156 | 295-305 | 170 | 138 | 160 | 312 |

### 2.2. Preprocessing

Task data were part of the “minimally preprocessed” release, which included gradient unwarping, fieldmap-based EPI distortion correction, brain-boundary-based registration of EPI to structural T1-weighted scan, non-linear registration, and intensity normalization [17]. Cortical data were mapped to a surface representation and utilized here. In addition, we regressed out 12 motion-related variables (6 translation parameters and their derivatives) and low frequency signal changes using the 3dDeconvolve program of the AFNI package [10] with the `ortvec` and `polort` options (the latter removed linear, quadratic, and cubic trends over the duration of individual runs. Resting-state also followed the so-called minimal pre-processing pipeline, in addition to denoising using ICA-FIX [34] and regressing out 12 motion-related variables, as provided with the data distribution. Cortical data were mapped to a surface representation. Preprocessing included minimal temporal filtering that essentially removed linear trends in the data. The ICA-FIX procedure removed “baed” components such as high frequency noise from the data. No further preprocessing was performed for resting-state data in the main text. In particular, band pass filtering is not included in HCP’s preprocessing because they believe it can potentially eliminate relevant information in resting-state data [6].

For the results in the main text, the global mean was not regressed from the data. In the supplemental material (Section S2), we repeated some analyses on resting-state data that included global signal regression as part of the preprocessing pipeline. Although there is no consensus in the field whether or not the global mean should be eliminated, some work has reported that removal strengthens the association between resting-state functional connectivity and behavior [20, 23].

### 2.3. Regions of interest and organization into subnetworks

For simplicity, we focused on cortical regions of interest (ROIs) only. We used the local-global Schaefer cortical parcellations that divide the cortex into 300 ROIs [32] (throughout the text, we refer to it as “whole-cortex”). A summary ROI-level time series was obtained by averaging signals within the region. We then used the Yeo 7-network parcellation to group the ROIs into 7 subnetworks known as `visual, somatomotor, dorsal attention, ventral attention, limbic, frontoparietal, and default mode` [39]. The number of ROIs within each of the subnetworks is provided in Table 2. The ROIs and the grouping into 7 networks is shown in Fig. S4. Some of the effects of varying the number of ROIs are described in the supplemental material (Section S3).

**Table 2:** Number of ROIs in each subnetwork. We used local-global Schaefer cortical parcellations that divide the cortex into 300 ROIs [32].

| Subnetwork | Visual | SomatoMotor | Dorsal Attention | Ventral Attention | Limbic | FrontoParietal | Default |
| --- | --- | --- | --- | --- | --- | --- | --- |
| Number of ROIs | 47 | 57 | 34 | 34 | 20 | 40 | 68 |

### 2.4. Functional connectivity

Functional connectivity was computed by Pearson correlating time series data between every pair of ROIs, resulting in 300 × 300 FC matrices. A symmetric matrix *S* that satisfies *y*′*S y* ≥ 0 (where *y*′ is the transpose of *y*) for any non-zero vector *y* is said to be positive semidefinite and has eigenvalues greater than or equal to zero. After normalizing the time series of each ROI to unit variance, let *x*_*t*_ = (*x*_*t*,1_, *x*_*t*,2_, …, *x*_*t*,300_) be the vector of activations of all ROIs at time *t* for *t* = 1, 2, …, *T*. If we denote the mean across time as 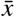, the covariance matrix is given by

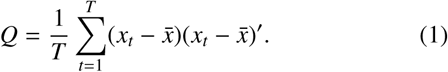

Note that the (*i, j*) entry of *Q* is simply the Pearson correlation coefficient between the time series of regions *i* and *j*. For any non-zero vector *y* of dimension 300,

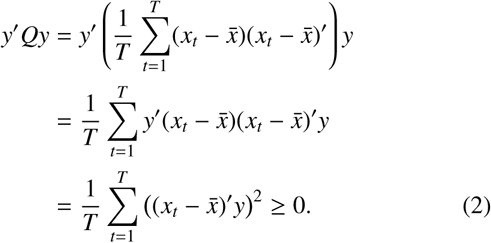

Thus, covariance matrices are positive semidefinite [7].

If *Q*_1_ and *Q*_2_ are two FC matrices, it can be easily shown following the steps above that *αQ*_1_ + *βQ*_2_ is also positive semidefinite for *α, β* > 0. Thus, the set of all positive semidefinite matrices lie on a cone referred to as the *positive semidefinite cone* [7].

### 2.5. Geometry of functional connectivity matrices

Pearson correlation is often used to characterize the similarity of FC matrices. However, as correlation matrices lie on a non-linear space, a natural approach is to compute *geodesic distances* between FC matrices to quantify their distance. The geodesic distance between two points on the positive semidefinite cone, and thus between two FC matrices *Q*_1_ and *Q*_2_, is the shortest path between them along the manifold [28]. There exists only one geodesic path joining two such points.

For two functional connectivity matrices, their geodesic distance can be computed as proposed in [28]:

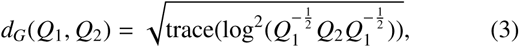

where the matrix log operator is used here. Note that this definition assumes that the matrix *Q*_1_ is invertible; when this was not the case the identity matrix, *I*, was added as a perturbation matrix to both *Q*_1_ and *Q*_2_ to ensure that all eigenvalues were greater than 0 (see Section S4). For matrices *Q*_1_ and *Q*_2_ of size *n* × *n* (here, *n* = 300 ROIs), if 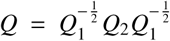, and *λ*_*i*_ for *i* = 1 to *n* are the *n* eigenvalues ≥ 0 of *Q*, the geodesic distance is simply (see https://github.com/makto-toruk/FC_geodesic for code)

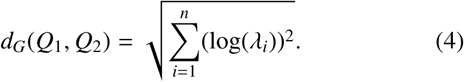

From (4), it is clear that *d*_*G*_ ≥ 0. In addition, *d*_*G*_ = 0 implies λ_*i*_ = 1 (i.e, *Q*_1_ = *Q*_2_), and vice versa. To verify that the geodesic distance is symmetric, note that *d*_*G*_(*Q*_1_, *Q*_2_) = *d*_*G*_(*Q, I*) (using Eq. 3). By the property of the log operator, *d*_*G*_(*Q, I*) = *d*_*G*_(*I, Q*) since log^2^(*Q*^−1^) = log^2^(*Q*). We refer the interested reader to [16] for a proof of the triangular inequality for 2 × 2 matrices. Thus, the geodesic distance applied to matrices meets the criteria of a *metric*.

If *q*_1_ and *q*_2_ are vectors obtained by stacking the columns of *Q*_1_ and *Q*_2_, respectively, Pearson dissimilarity between the two matrices is defined as

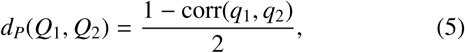

where the corr function is the Pearson correlation coefficient. Pearson dissimilarity ranges between 0 and 1 and is *not* a formal metric because it does not satisfy the triangular inequality [36]. The units for geodesic distance and Pearson dissimilarity are arbitrary and thus not comparable across these measures.

### 2.6. Participant identification

Identification involves mapping an unknown participant’s data to one of the participants in the database. Since each task in the HCP data contains 2 runs for every participant, we used one run as training data (that is, to form the database) and the other run for testing. Identification was performed on each condition (resting-state or task) separately.

Participant identification is equivalent to *N*-class classification where the objective is to label an individual’s FC matrix in the test data to one of the *N* participants in the training data. To do so, we used a 1-Nearest Neighbor approach [15]: An FC matrix in the test data is labeled with the participant identity of the FC that is most similar to it in the training data. Suppose 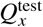 is an unknown participant’s FC matrix. Then

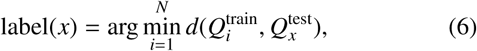

where 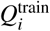 is the *i*th participant’s FC matrix in the training data and *d*(·,·) is a distance or similarity measure. Here we compare the use of a geodesic distance metric to a Pearson dissimilarity measure.

#### 2.6.1. Identification accuracy

Participant identification was performed using the first run as training data and the second run as testing data. For the *N* participants in the testing data, accuracy was defined as

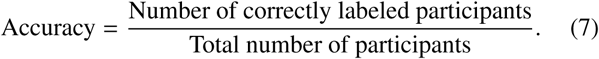

Then, the roles of the training and testing data were reversed and accuracy was computed again. The reported identification accuracy was the mean of the two accuracy values.

### 2.7. Bootstrapping

For participant identification statistics, one must confront the non-independence between participants in the sample. Consider the following case. If two participants’ FC matrices *Q*_*A*_ and *Q*_*B*_ are close to each other, *B* might be mislabeled as *A*. However, if *A* was not in the training database, it is conceivable that *B* would have been labeled correctly. Therefore, the *entire group* must be considered as the unit of interest; it is the group that determines if identification performance will be poor or good. In our study, we used data from *N* = 100 participants in the age range of 22 − 35 years, but demographic factors such as age and mental health status can potentially play an important role in identification performance.

A convenient procedure to assess variability in identification performance is to use bootstrap resamples, with each resample comprising random draws with replacement of the urn containing the group of participants. Thus, a bootstrap resample is a proxy for a group of participants, and variability can be quantified by resampling it a large number of times.

More precisely, suppose a dataset of size *N* for a run is denoted by 𝒟. Let 0 ≤ *f*_*P*_(𝒟) ≤ 1 and 0 ≤ *f*_*G*_(𝒟) ≤ 1 be the participant identification accuracy obtained using Pearson dissimilarity and the geodesic distance, respectively. Let ℛ_*j*_ be a dataset also of size *N* obtained by resampling 𝒟, with replacement, *N* times. Thus, ℛ_*j*_ is a bootstrap resample of 𝒟 and may contain duplicate entries. The accuracy difference on this bootstrap resample is given by d(ℛ_*j*_) = *f*_*G*_(ℛ_*j*_) − *f*_*P*_(ℛ_*j*_). Such difference score is computed for *M* = 1000 bootstrap resamples ℛ_1_, ℛ _2_, …, ℛ_*M*_ and the *mean* difference score, 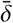, is computed. This process (based on *M* resamples) provides exactly one mean difference score. The question of interest is as follows: How are *mean* difference scores distributed? Note that this parallels the question of the distribution of the sample mean in the setting of the standard Central Limit Theorem. In our case, the distribution of *mean difference scores* is of interest. Since the object of interest is the *mean* difference score, the procedure to determine a specific 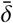 is repeated *B* = 1000 times, resulting in 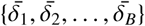 (that is, *B* mean differences).

Although the number of resamples, *M* × *B*, is large, the distance matrix of size *N* × *N* (between each subject’s test-FC to all subjects’ train-FC) is calculated only once making the bootstrapping procedure computationally feasible.

Reported *p*-values were computed as follows. Because accuracy differences are percentages, we initially applied a standard Fischer-*z* transformation to 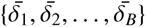 so that their distribution would be approximately normal. To test the null hypothesis 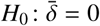, a one-sample *t*-test was then used.

#### 2.7.1. Evaluating shorter data segments

To understand the effect of the length (or the number of frames) of the run, we truncated runs to smaller segments. For a particular segment length, 50 segments were obtained each of which had a unique, randomly-chosen starting point in the run. The objective was to pick several segments of the same length without favoring those that started at the beginning of the scan. For each segment, 1000 bootstrap iterations were used to obtain a mean accuracy score.

### 2.8. Multidimensional scaling

Naturally, visualizing distances between FC matrices is not straightforward given their high dimensionality. Here, we used *non-metric multidimensional scaling* to visualize distances in three dimensions [21]. Whereas standard multidimensional scaling computes the Euclidean distance between the high-dimensional vectors of interest, non-metric multidimensional scaling takes as input any *dissimilary matrix* of the form

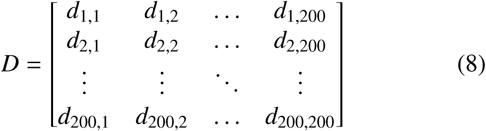

where *d*_*i,j*_ is the “dissimilarity” between the FC matrices *i* and *j* (the dimensionality of the matrix is 200 since we consider a test-FC and a train-FC for each of the *N* = 100 participants). Here, either geodesic distance or Pearson dissimilarity was used. Given *D*, non-metric multidimensional scaling finds a set of ℝ^3^ vectors such that the Euclidean distance between these vectors preserves, to the extent possible, the high-dimensional distances:

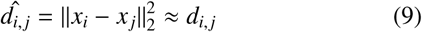

where the vectors *x* are low dimensional. Thus, if *d*_*i, j*_ = *d*(*Q*_*i*_, *Q*_*j*_) is the distance between two FC matrices *Q*_*i*_ and *Q*_*j*_, and 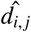 is the distance in the lower-dimensional representation, the output (set of points) is produced by minimizing the *stress* function:

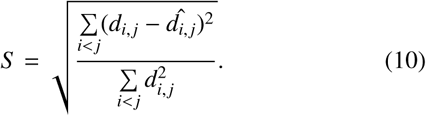

The optimal distances, 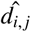, are obtained using a gradient descent approach that minimizes the stress. The MATLAB 2018a [24] implementation of *mdscale* with 1000 gradient descent iterations was used. Multidimensional scaling produces low dimensional representatives, *x*’s, for high dimensional FCs such that the Euclidean distances between *x*’s approximate the measured relationships (Pearson dissimilarity or geodesic distance) between their high-dimensional counterparts. Given that the two measures have arbitrary units, so do their estimates in low dimensions.

Note that the objective of using non-metric multidimensional scaling was to represent in a more intuitive manner the relationships between high-dimensional functional connectivity matrices. Thus, points in the lower-dimensional representation no longer lie on the positive semidefinite c one a nd *closeness* should be interpreted in the Euclidean sense (two points are close if their Euclidean distance is small). The visualizations, approximate as they are, are only provided to aid understanding, and are not part of the procedure to determine identification accuracy.

### 2.9. Note on p-values

As discussed by many others recently, we do not view “statistical significance” dichotomous thresholds (for example, *p* < 0.05) as the ultimate criterion in deciding whether a result is “real” or not ([4, 25]). In any case, understanding variability and the unlikeliness of a result provides some information. Given that we compare geodesic distance to Pearson dissimilarity across conditions and other parameters, some form of correction for multiple comparisons is opportune. Accordingly, we provide the uncorrected *p*-value as well as the Bonferroni-corrected *α* level (which we call the “reference *α*”) so that the reader can further gauge the “strength” of the finding. Again, we do not advocate using the Bonferroni-corrected *α* in a dichotomous fashion, but provide it as an additional “reference” point for the reader.

## 3. Results

### 3.1. Motivation behind geodesic distance

We motivate the geodesic distance with simple examples from the space of 2 × 2 FC matrices. Since FC matrices are symmetric and positive semidefinite, they take the form

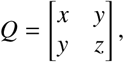

and satisfy *x* ≥ 0, *y* ≥ 0 and *xy* − *z*^2^ ≥ 0. Since the matrices have only three unique entries, all points that satisfy these equations can be plotted in three dimensions in Euclidean space, and form a *positive semidefinite cone* (Fig. 1B).

In the first example, we considered three points on the cone (i.e., three 2 × 2 FC matrices) ‘*a*’, ‘*b*’ and ‘*c*’ such that ‘*b*’ and ‘*c*’ are equidistant from ‘*a*’ in terms of the Euclidean distance (Fig. 2A). If a tangent surface to the cone is drawn at ‘*a*’, the point ‘*c*’ is much closer to the tangent surface than ‘*b*’. Thus, the geodesic distance between ‘*a*’ and ‘*b*’ is larger than that between ‘*a*’ and ‘*c*’ (Fig. 2B). In this case, Pearson dissimilarity is capable of distinguishing the two distances.

**Fig. 2:**
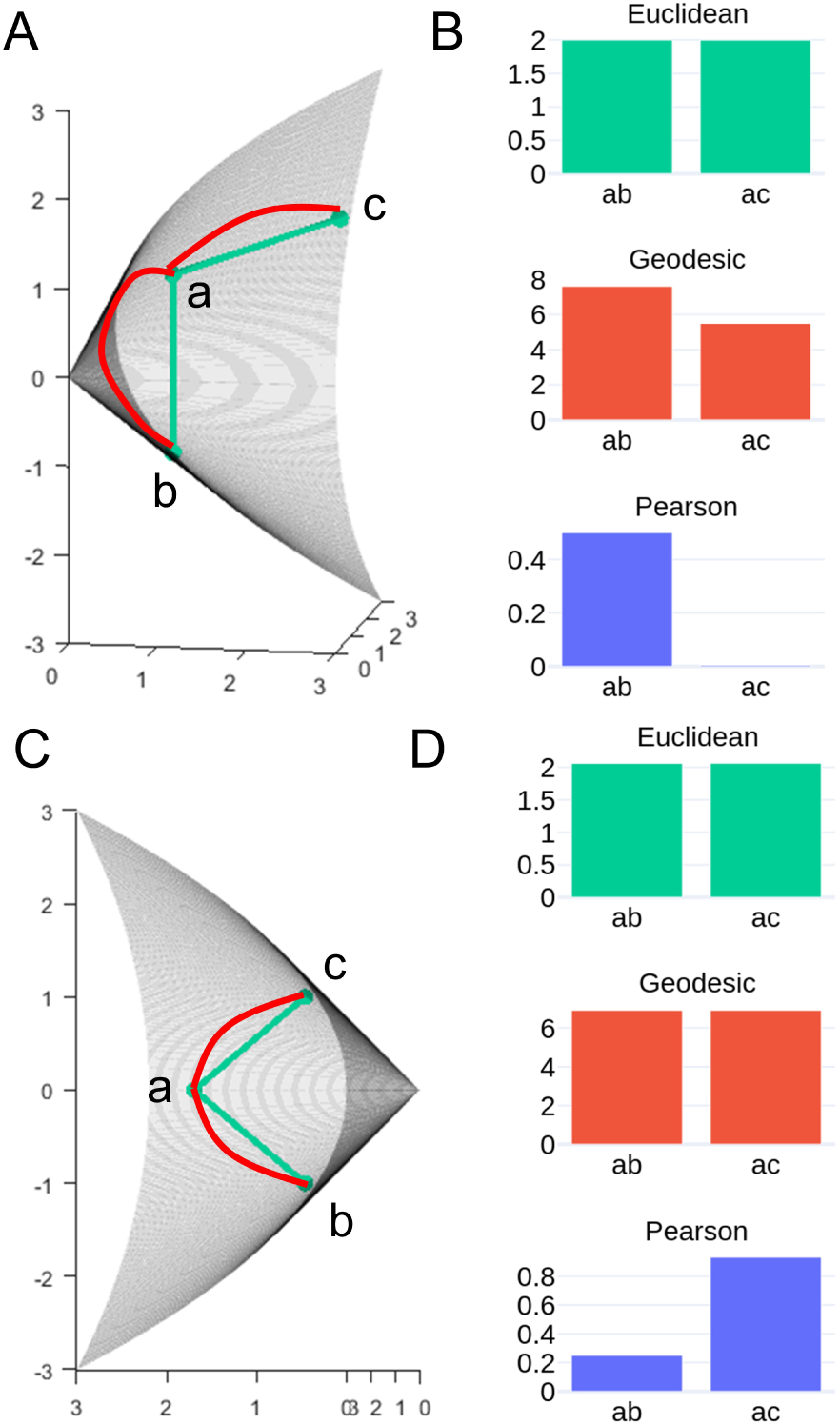
Motivating functional connectivity geometry. (A) Identical Euclidean distance does not imply identical geodesic distance. (C) Identical geodesic distance can yield very different Pearson dissimilarity. (B, D) Comparison of distances/dissimilarity *ab* and *ac* in (A) and (C), respectively. Distances/dissimilarity cannot be compared across measures because their units are arbitrary.

To motivate why Pearson dissimilarity is problematic, consider that the Pearson correlation between two vectors is equivalent to the cosine of the angle between them after they have been “centered” individually (that is, the mean of each vector is subtracted from it) and normalized. Indeed, the computation of Pearson correlation eliminates the contribution of the signal mean, as can be readily seen in the following equation:

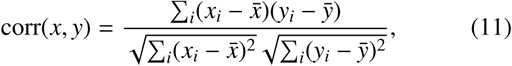

where *x* and *y* are vectors. For FC matrices, such centering which is implicit in Pearson correlation alters the eigenvalues and the positive semidefiniteness of the matrix. Since the eigenvalues are the basis for computing geodesic distances, we see that Pearson correlation in fact distorts the evaluation of similarity between connectivity matrices (relative to what is estimated with the geodesic distance). However, while estimating an individual’s FC matrix, mean centering does not affect positive semidefiniteness, as shown in Eq. (2).

In a second illustrative example (Fig. 2c), we consider three points ‘*a*’, ‘*b*’ and ‘*c*’ on the cone such that ‘*b*’ and ‘*c*’ are symmetrically on either side of ‘*a*’. By symmetry, ‘*a*’ is equidistant from ‘*b*’ and ‘*c*’ in terms of both the Euclidean distance and geodesic distance. However, Pearson dissimilarity between the two sets of points can be quite distinct. Suppose *O* is the origin and ∠*aOb* = ∠*aOc* (where ∠ is the angle subtended between ‘*a*’ and ‘*b*’). Since Pearson correlation mean centers the vectors ‘*a*’ and ‘*b*’, the correlation is related to mean-centered vector angles that can be quite different from the original ones (Fig. 2d). In other words, if ‘*<u>a</u>*’, ‘*<u>b</u>*’, and ‘*<u>c</u>*’ are vectors obtained by centering ‘*a*’, ‘*b*’ and ‘*c*’, in most cases ∠*<u>a</u>O<u>b</u>* * ∠*<u>a</u>O<u>c</u>*. The upshot is that measures of similarity based on Pearson correlation do not correspond to actual distances between functional connectivity matrices.

### 3.2. Geodesic distance and participant identification

Participant identification (*N* = 100) was performed on each condition (resting-state and tasks) using two measures: geodesic distance and Pearson dissimilarity (Methods 2.6). FC matrices obtained from one run were used as training data and matrices from the second run as testing data. Identification accuracy for each condition is shown in Fig. 3 (accuracy based on chance would be 1%).

**Fig. 3:**
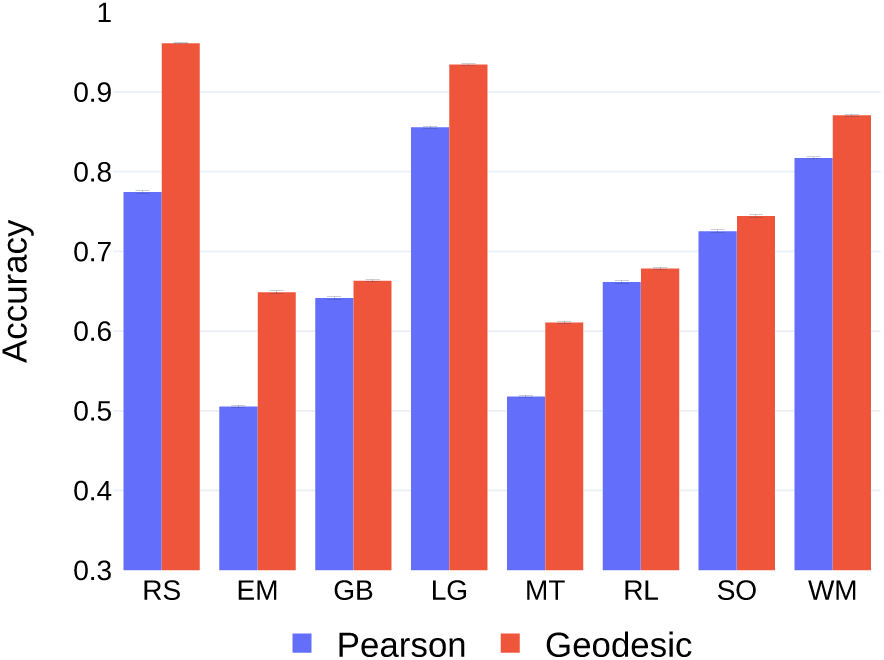
Participant identification for the eight conditions using the geodesic distance and Pearson dissimilarity. Training and testing data were from the same condition. Accuracy improved using the geodesic distance on each condition. Error bars indicate standard error of the mean across bootstrap iterations. Abbreviations: EM, emotion processing; GB, gambling; LG, language; MT, motor; RL, relational processing; RS, resting-state; SO, social cognition; WM, working memory.

To assess the robustness of the results and for statistical comparisons between the two measures, identification was performed on bootstrap resamples. For each bootstrap resample, the difference between accuracy using geodesic distance and Pearson dissimilarity was computed. A one-sample two-tailed *t*-test was then used to assess the null hypothesis that the difference distribution had zero mean (Methods 2.7). For each condition, using the geodesic distance improved identification accuracy over Pearson dissimilarity (*p* < 10^−6^ for all tasks; reference *α* = 0.05/8 = 0.00625 given 8 conditions; Fig. S6). The mean improvement using geodesic distance was around 8%, ranging from 2% (*relational processing*) to as much as 19% (*resting-state*). For *resting-state* and the *language* conditions, the accuracy obtained using the geodesic distance was very hight and close to 95%.

Finn et al. [15] reported a mean accuracy of 93.65% on *resting-state* data using Pearson dissimilarity, which is considerably higher than the 77.5% we obtained. Given that in the HCP dataset four runs of *resting-state* data are available per participant (collected over separate days), they averaged the FC matrices obtained obtained during the same day into a single FC matrix^1^. By including this averaging procedure, we replicated their findings more closely and obtained an accuracy of 91% using Pearson dissimilarity. Using geodesic distance, accuracy increased to 98%. However, since conditions other than *resting-state* contained only two runs, we did not use the averaging procedure on the four runs of resting-state data in the remainder of our work.

### 3.3. Low-dimensional visualization of functional connectivity matrices

Since FC matrices are high dimensional, multidimensional scaling was used to visualize the distances between them in three dimensions (Fig. 4). The goal of using multidimensional scaling was to represent in a more intuitive manner the relationships between high-dimensional FC matrices. Accordingly, points in the lower-dimensional representation should be interpreted in the Euclidean sense (two points are close if their Euclidean distance is small). But note that the visualizations are approximate only, and provided to aid understanding (they are not part of the procedure to determine identification accuracy).

**Fig. 4:**
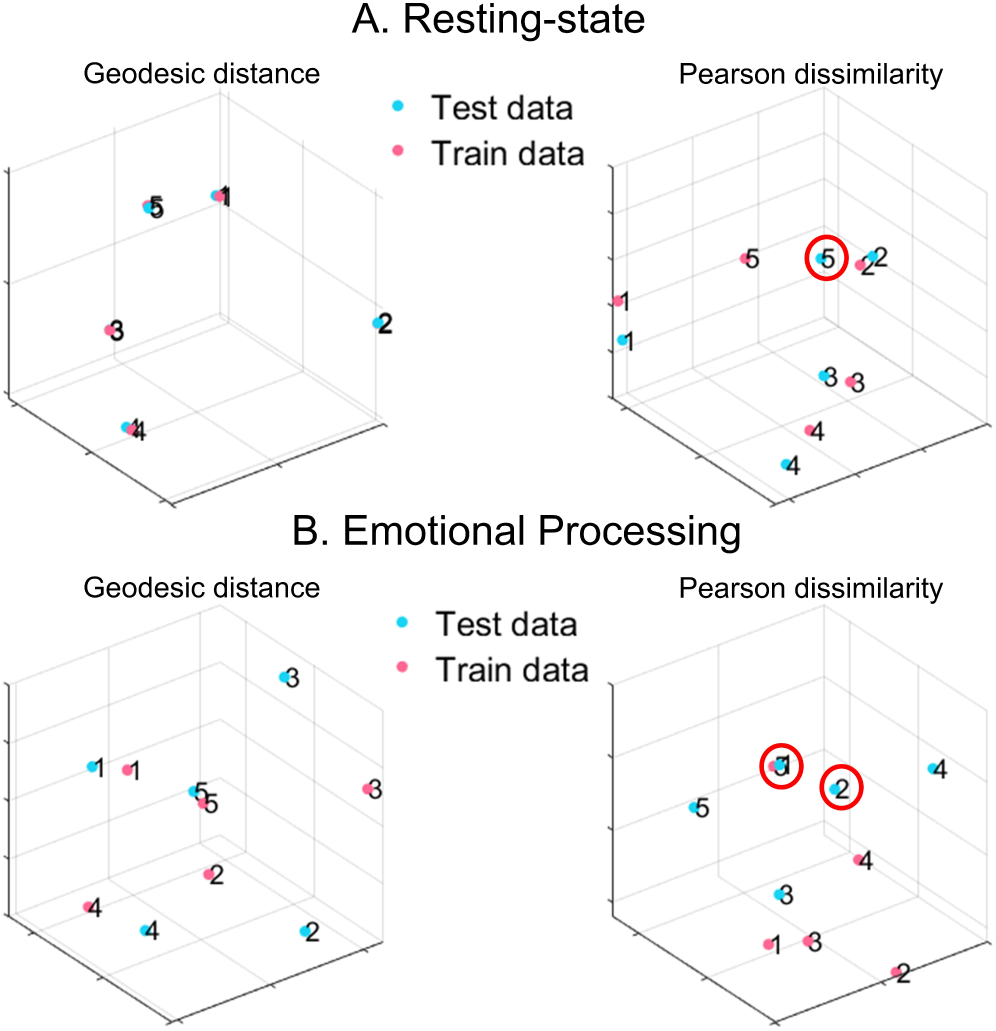
Visualization of geodesic distance and Pearson dissimilarity. Distance/similarity between high-dimensional functional connectivity matrices (300 × 300) was visualized in three dimensions using non-metric multi-dimensional scaling. Training data (blue) and testing data (pink) were selected from five random participants (numbers 1-5). Mislabeled participants are encircled in red. (A) *Resting-state*. (B) *Emotional processing* task. For *resting-state*, within-participant geodesic distances were very small relative to between-participant distances in the lower-dimensional representation (when numbers labeling the participants overlapped, only one of them is visible). Online figures are available [37].

Within- and between-participant distances estimated in three dimensions were indicative of varying identification accuracy (obtained using high dimensional FC matrices) across conditions. For *resting-state*, FC matrices within-participant geodesic distances between training and testing were very small, whereas distances between different participants were considerably larger, consistent with the high identification accuracy. Visualization of Pearson dissimilarity revealed similar characteristics, but the ratios of within- to between-participant distances were not as large. In fact, using Pearson dissimilarity resulted in participant 5 being mislabeled as participant 2, for example.

For the *emotional processing* task, within-participant distances were *not* much smaller than between-participant distances even for the geodesic distance consistent with the lower accuracy on this task. However, all participants in the randomly chosen subset were still labeled correctly. Using Pearson dissimilarity, two participants were mislabeled. In general, using the geodesic distance resulted in more favorable ratios of within- to between-participant FC distances.

### 3.4. Identification accuracy and time course length: resting-state data

Since the length of the time course plays a key role in the quality of the estimate of the FC matrix [22, 40], we sought to characterize its effect on participant identification. Because *resting-state* data had the longest time course (1200 TRs), shorter segments varying from 100 to 1100 TRs (in steps of 100) were extracted. Accuracy improved with length for both measures (Fig. 5). Accuracy using the geodesic distance was higher than Pearson dissimilarity for segment lengths greater than 200 TRs (*p* < 10^−4^; reference *α* = 0.05/11 = 0.0045 given 11 segment lengths; Fig. S7). For segment length of 100 TRs, accuracy using geodesic distance was still higher than Pearson dissimilarity (but *p* = 0.051). Notably, the geodesic distance, with segment lengths as short as 300 TRs, outperformed the best accuracy using Pearson dissimilarity which was obtained with the full time course (four times more data; *p* < 10^−4^; reference *α* = 0.05/11 = 0.0045 given 11 segment lengths).

**Fig. 5:**
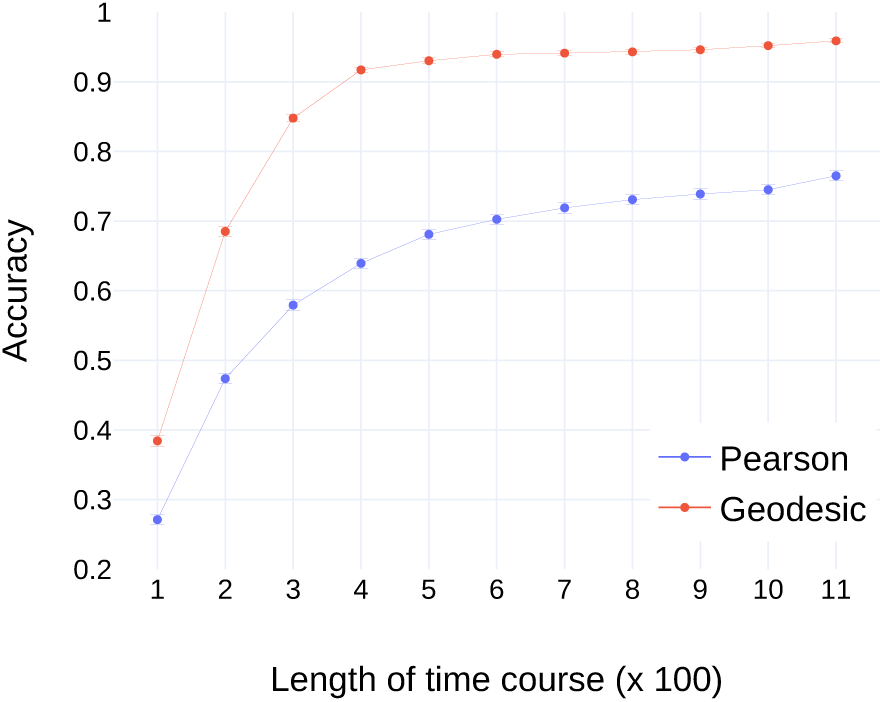
Participant identification accuracy as a function of segment length for *resting-state* data. Accuracy using geodesic distance exceeded Pearson dissimilarity at each segment length (see text). Error bars indicate standard error of the mean across bootstrap iterations.

### 3.5. Identification accuracy and time course length: task data

Although accuracy increased with segment length for *resting state*, length did not predict performance straightforwardly (Fig. 6A). In particular, *working memory* and *language* tasks had comparable time course lengths, but identification accuracy differed by as much as 10%. To probe this issue further, runs were trimmed so that they all had the same duration (138 TRs, which was the length of the shortest task; for conditions with more data, this target length was obtained by deleting time points at the beginning and end of the data segment, thereby retaining the middle part).

**Fig. 6:**
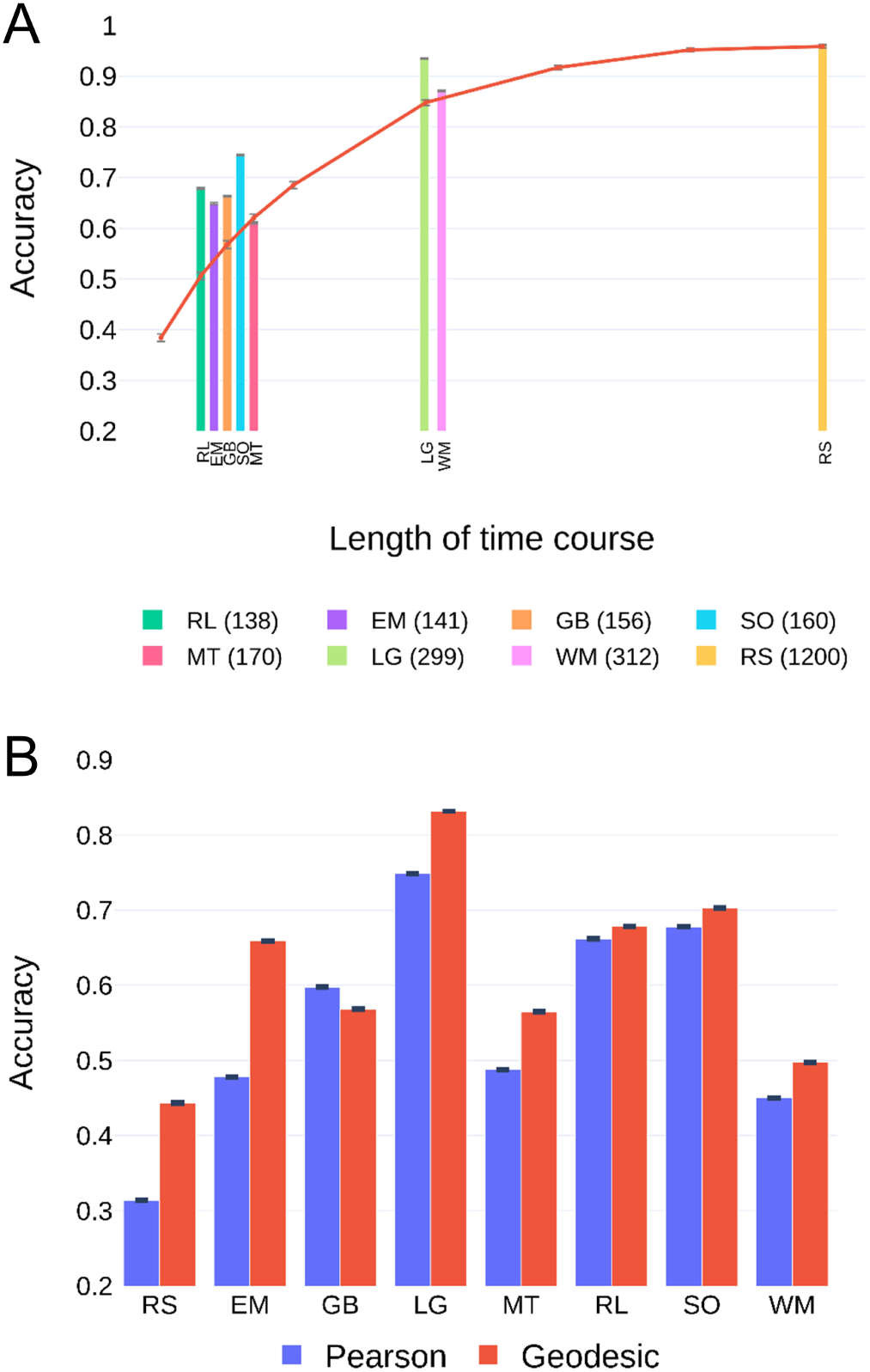
Participant identification and time course length. (A) Accuracy based on geodesic distance for *resting-state* and task conditions (time course length in TRs in the inset). The red curve shows the accuracy for *resting-state* data trimmed to segment lengths shorter and longer than those of task data (lengths from left to right: 100, 125, 145, 170, 200, 300, 600, 900, and 1200 TRs). (B) Accuracy when data was trimmed such that all conditions had the same time course length (138 TRs). Error bars indicate standard error of the mean across bootstrap iterations. Abbreviations: EM, emotion processing; GB, gambling; LG, language; MT, motor; RL, relational processing; RS, resting-state; SO, social cognition; WM, working memory.

With time course length equated, accuracy still varied considerably across tasks (Fig. 6B). Accuracy obtained using the geodesic distance exceeded that of Pearson dissimilarity for all conditions except the *gambling* task (*p* = 1 for *gambling, p* < 10^−4^ for all other tasks; reference *α* = 0.05/8 = 0.00625 given 8 conditions; Fig S8). Notably, although *resting-state* had the highest identification accuracy when the entire time course was used, it had the lowest identification accuracy when length was equated across conditions.

### 3.6. Brain subnetworks and participant identification

Particular brain subnetworks are known to be engaged more prominently, as well as exhibit enhanced functional connectivity, during particular tasks [29]. To evaluate performance based on subsets of regions, ROIs were grouped into seven subnetworks (Methods 2.3). Was the best subnetwork for identification dependent on condition? Data for all conditions were trimmed so that they had the same length (138 TRs; the `limbic` subnetwork was excluded because identification accuracy was less than 10% across conditions).

Using geodesic distance improved the accuracy across most conditions for most subnetworks (Fig. 7A). In particular, for the `visual, dorsal attention, frontoparietal` and `default mode` subnetworks, accuracy was comparable to that obtained with the whole cortex. For example, the `default mode` subnetwork produced accuracy over 90% for the *language* task. The `frontoparietal` performance on *resting-state* and *emotion processing* was close to 80%. Further inspection of Fig. 7A revealed additional features of condition/subnetwork combinations. For example, the `visual` subnetwork was not very suitable for identification based on *resting-state* data. Not surprisingly, the `default mode` subnetwork performed well with *resting-state* data. Interestingly, the `frontoparietal` subnetwork performed nearly as well with *resting-state* data, too. These two subnetworks obtained even higher identification accuracy during the *language* task.

**Fig. 7:**
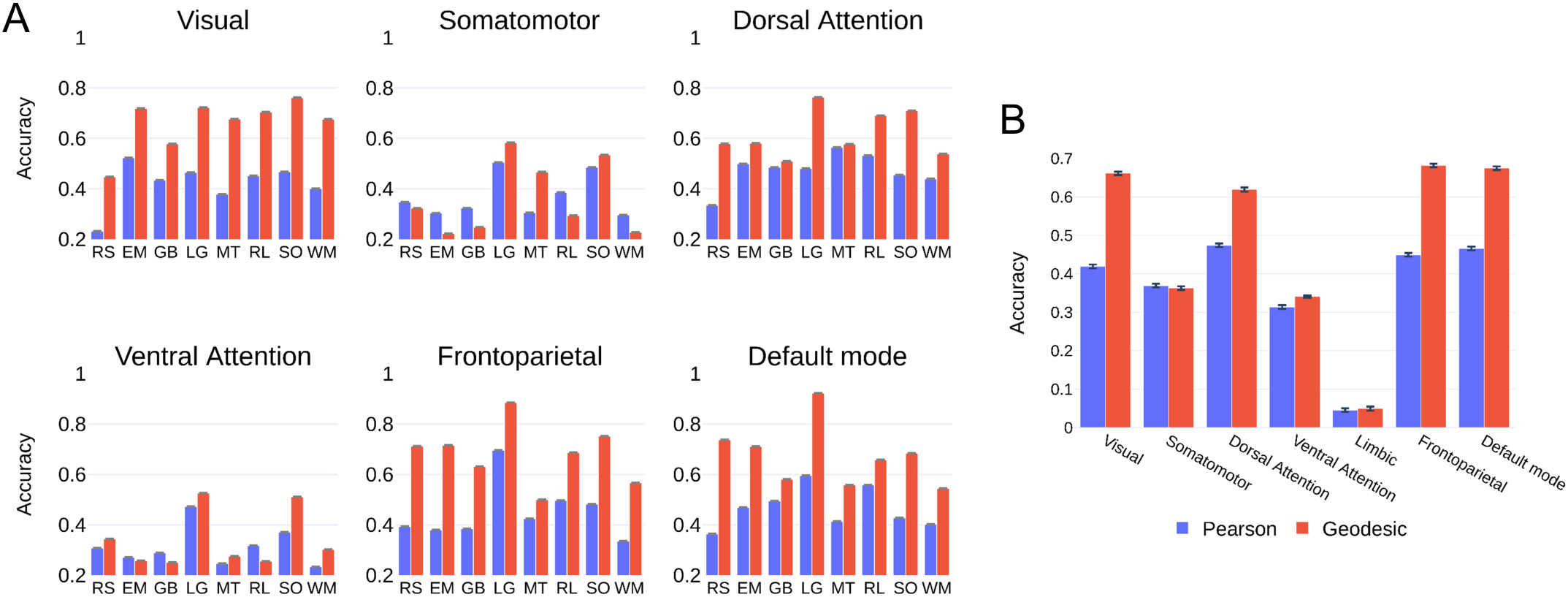
(A) Participant identification accuracy using subnetworks. Runs were trimmed such that all conditions had the same time course length. Some subnetworks were more suitable than others for identifying individual differences. The use of geodesic distance showed considerable improvements in accuracy for most subnetworks. (B) Across subnetworks, average participant identification accuracy is displayed. The geodesic distance substantially improved identification accuracy. Error bars indicate standard error of mean across bootstrap iterations. Abbreviations: EM, emotion processing; GB, gambling; LG, language; MT, motor; RL, relational processing; RS, resting-state; SO, social cognition; WM, working memory.

To further evaluate performance of subnetworks, identification accuracy was averaged across conditions (Fig. 7B). By using the geodesic distance, accuracy improved substantially, with several subnetworks improving by over 20%. Except for the `somatomotor` subnetwork, using the geodesic distance resulted in improved performance (*p* = 0.996 for `somatomotor`, *p* < 10^−5^ for all other subnetworks; reference *α* = 0.05/7 = 0.0071 given 7 subnetworks; see Fig. S9 for bootstrap distributions). The highest mean accuracies were observed in the `visual, dorsal attention, frontoparietal`, and `default mode` networks for both geodesic and Pearson measures, indicating that some subnetworks are more suitable than others for participant identification.

Fig. 8 displays geodesic identification accuracy for each condition as a function of subnetwork size. Whereas the smallest subnetwork (`limbic`) performed poorly for all conditions, accuracy did not always increase with size. For example, the `dorsal attention` and `ventral attention` subnetworks have the same size, but the former produced considerably higher accuracy on each condition (*p* < 10^−12^ for all conditions; reference *α* = 0.05/8 = 0.00625 given the 8 conditions; see Fig. S10 for bootstrap distributions). Across conditions, the `dorsal attention` improved over the same-sized `ventral attention` by over 20%. Of note, the `somatomotor` subnetwork was larger than all but the `default mode` subnetwork, but it produced relatively low identification accuracy; at the same time, the largest subnetwork (`default mode`), was associated with consistently high accuracy across conditions. Finally, no single subnetwork exhibited the highest accuracy for all conditions. In fact, performance varied across conditions, but also varied in particular ways across subnetworks for each condition. Notably, the `visual`, textttdorsal attention, `frontoparietal`, and `default mode` subnetworks performed consistently well. Similar trends were observed for the Pearson dissimilarity measure but overall accuracy levels were lower (Fig. S11).

**Fig. 8:**
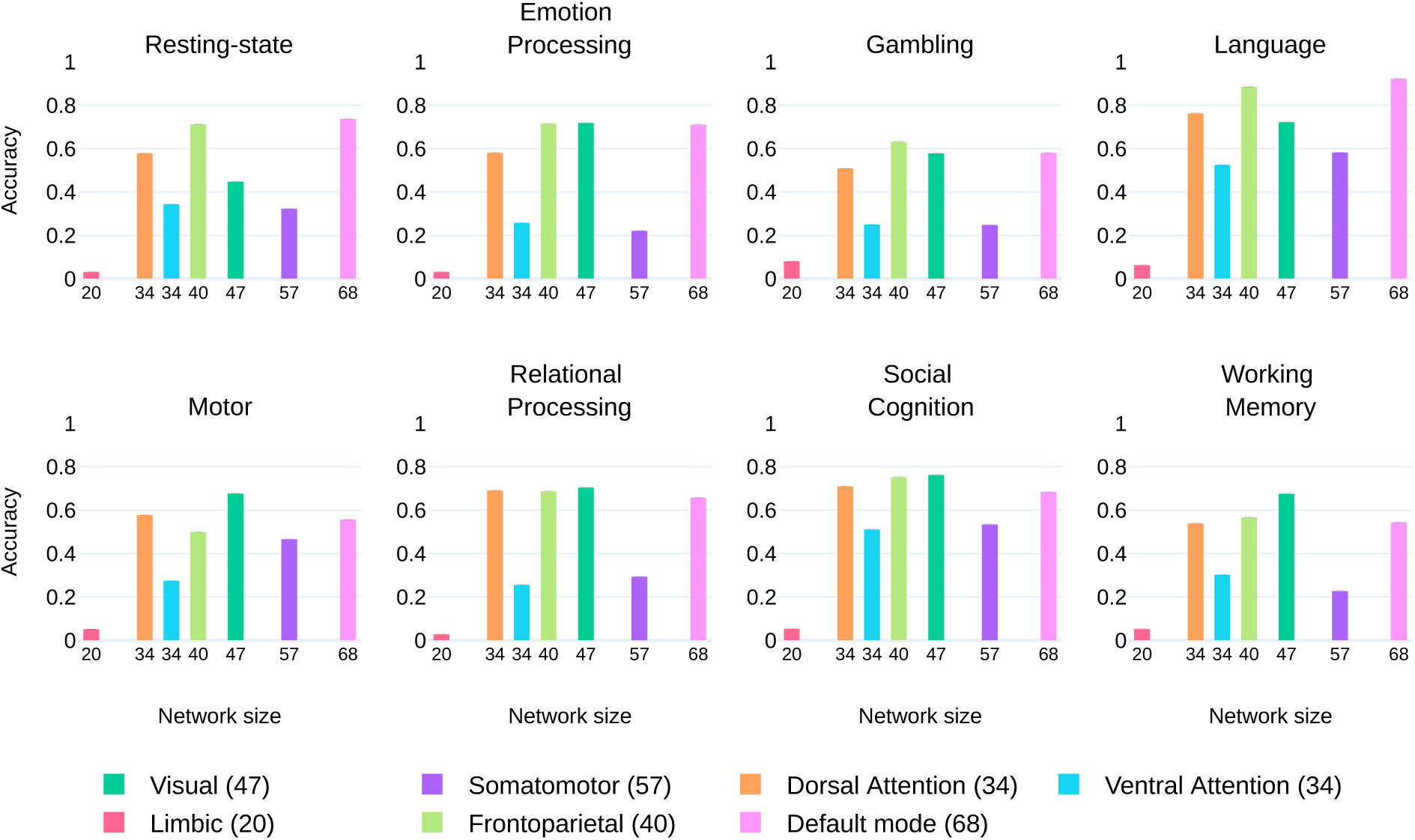
Participant identification accuracy plotted against subnetwork size for each condition (geodesic distance). The size of the subnetwork (the number of ROIs) is also indicated in the inset. The error bars represent standard error of the mean across bootstrap iterations.

### 3.7. Combining subnetworks improved identification accuracy

As described, subnetworks had comparable (and sometimes higher) identification accuracy than whole-cortex performance, although subnetworks were associated with much smaller matrices, of course. Could particular subnetworks be combined to further improve identification? We tested this possibility by targeting two subnetworks that exhibited high performance overall, namely `frontoparietal` and `default mode` (see Fig. 7B). The combined network included all within-network functional connections of course, but also all between-network links (for example, a functional connection between a region of the `frontoparietal` network and a region of the `default mode` network). Time course length was equated for all conditions as in Section 3.5. Accuracy using geodesic distance was superior to Pearson dissimilarity (Fig. 9; *p* < 10^−15^ for all conditions; reference *α* = 0.05/8 = 0.00625 given 8 conditions; Fig. S12).

**Fig. 9:**
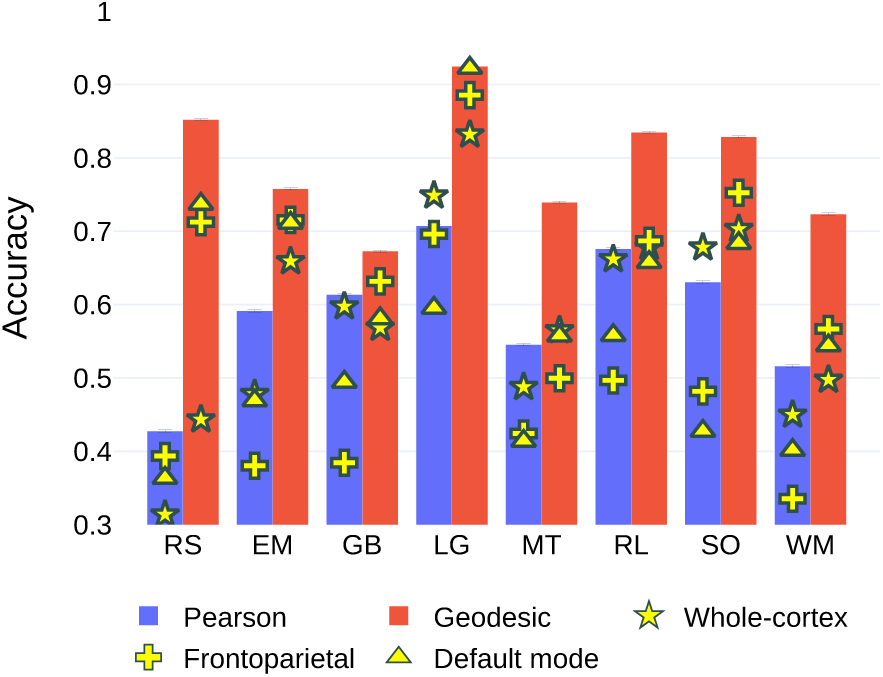
Participant identification accuracy by combining subnetworks. For the geodesic distance, the `frontoparietal` (`subnet1`) and `default mode` (`subnet2`) subnetworks were combined. For the Pearson dissimilarity measure, the `dorsal attention` (`subnet1`) and `default mode` (`subnet2`) subnetworks were combined (the top two subnetworks based on mean accuracy across conditions for this measure). Abbreviations: EM, emotion processing; GB, gambling; LG, language; MT, motor; RL, relational processing; RS, resting-state; SO, social cognition; WM, working memory.

Using geodesic distance, the combined subnetwork also outperformed both the individual subnetworks on all conditions except the *language* task (*p* = 0.24 for the *language* task, *p* < 10^−12^ for all other conditions; *α* = 0.05/16 = 0.003125 given 8 conditions and comparisons with two subnetworks; see Fig. S13-S14 for bootstrap distributions). In addition, for the geodesic distance, the combined subnetworks exhibited higher accuracy than whole-cortex FC matrices (*p* < 10^−12^ for all conditions; *α* = 0.05/8 = 0.00625 given 8 conditions; see Fig. S15 for bootstrap distributions) although the number of ROIs in the combined subnetwork (108) was nearly a third as those in the cortex (300). Clearly, the improvement in accuracy was not a simple consequence of increased size, but resulted from improved identity characterization.

To understand whether addition of other subnetworks to the combined network further improved accuracy, we performed identification using combinations of the seven networks taken two, three, and four at a time. The maximum identification accuracy across all combinations of subnetworks is displayed against the number of combined subnetworks in Fig. 10A. The minimum identification accuracy across the combinations of subnetworks is also indicated. For all conditions, accuracy initially increased as more subnetworks were considered but then decreased. Performance peaked at 2 or 3 subnetworks for all conditions. Accuracy varied across the combinations of subnetworks (when the number of subnetworks was held constant), and the minimum value (shown in yellow) was less than half the maximum in most cases. In Fig. 10B, identification accuracy was averaged across conditions and displayed as a function of the number of combined subnetworks.

**Fig. 10:**
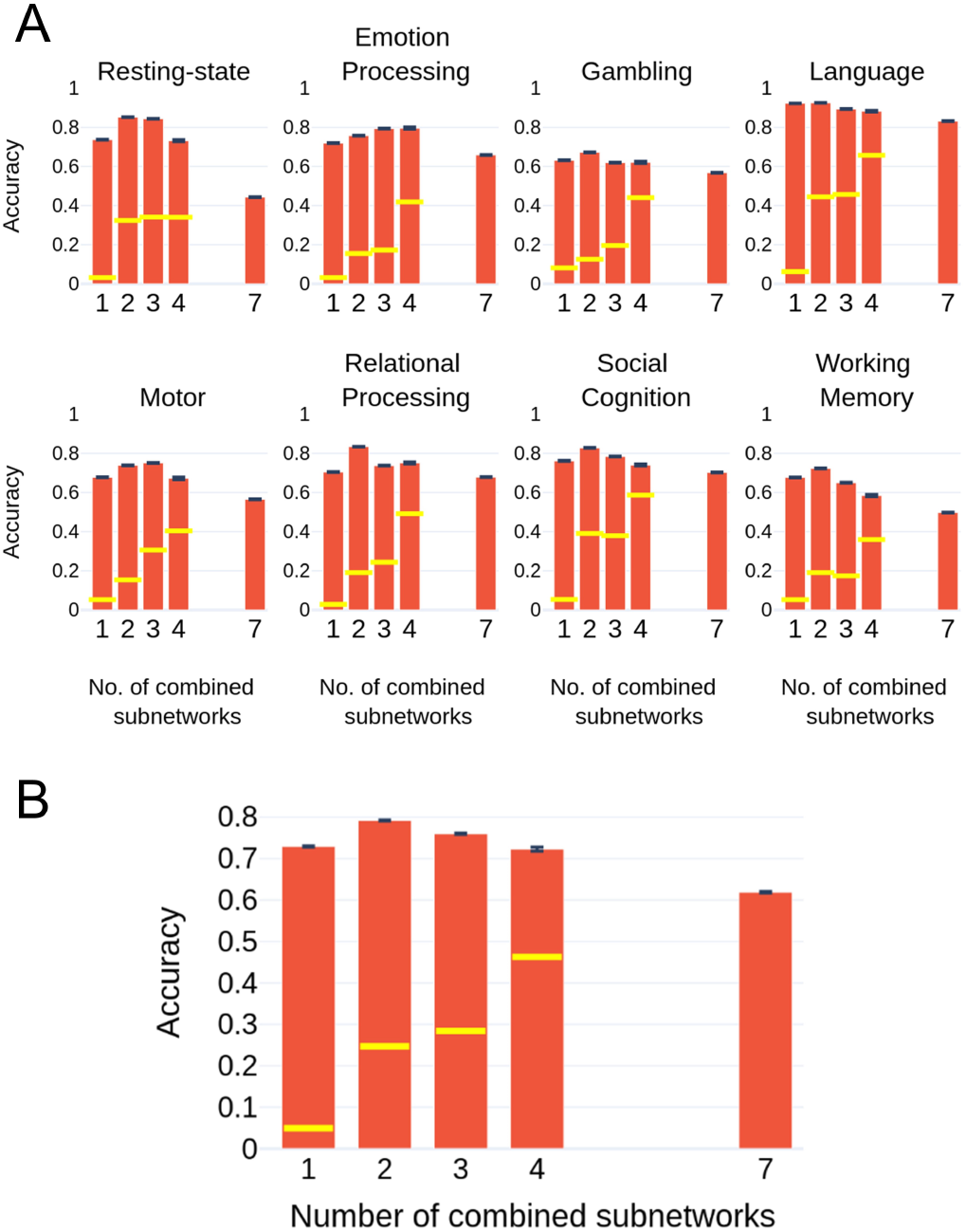
Combining up to seven subnetworks. (A) Participant identification accuracy using geodesic distance as a function of the number of subnetworks for each condition. For a particular condition and number of combined subnetworks (1, 2, 3, or 4), the maximum identification accuracy across all combinations of subnetworks is shown with the red bar (the minimum is indicated by the yellow line). Accuracy initially increased with the number of subnetworks but then decreased, and was lowest using whole-cortex FCs (i.e, number of combined subnetworks = 7). (B) Participant identification accuracy averaged across conditions is displayed against number of combined subnetworks.

### 3.8. Transfer of identifiability between conditions

In the previous sections, training and testing data were based on the same condition. Here, we sought to understand if participants could be identified if the training and testing data were obtained from different conditions; for example, identifying a participant performing a *working memory* task when the training used *resting-state* data. Time series length was not equated across conditions because our goal was to evaluate how transferable identity-related information was between pairs of conditions. Accordingly, we did not want to potentially degrade FC information by using shorter data segments. Identification was performed on the combined `default-plus-frontoparietal` network, which as discussed performed well across conditions (Fig. 9).

Results for both geodesic distance and Pearson dissimilary are displayed in Fig. 11. Whereas Pearson dissimilarity was useful in identifying participants when they performed the same task (within-conditions, diagonal entries), performance deteriorated when the training and test data originated from different tasks. Notably, across-condition identification was considerably higher with the geodesic distance, and this enhancement was rather striking when the training data was from *resting-state*, and to some extent based on the *language* and *working memory* tasks. For example, testing *working memory* data based on training with *resting-state* data yielded 94.6% accuracy, which intriguingly was even better than when training with *working memory* itself (accuracy: 92.9%, *p* < 10^−4^). On average, training with *resting-state* yielded 83.4% accuracy when testing on *other* conditions (see the “column mean” in Fig. 11). The present results indicate that the geometry of FC is especially important for across-task identification (see Discussion).

**Fig. 11:**
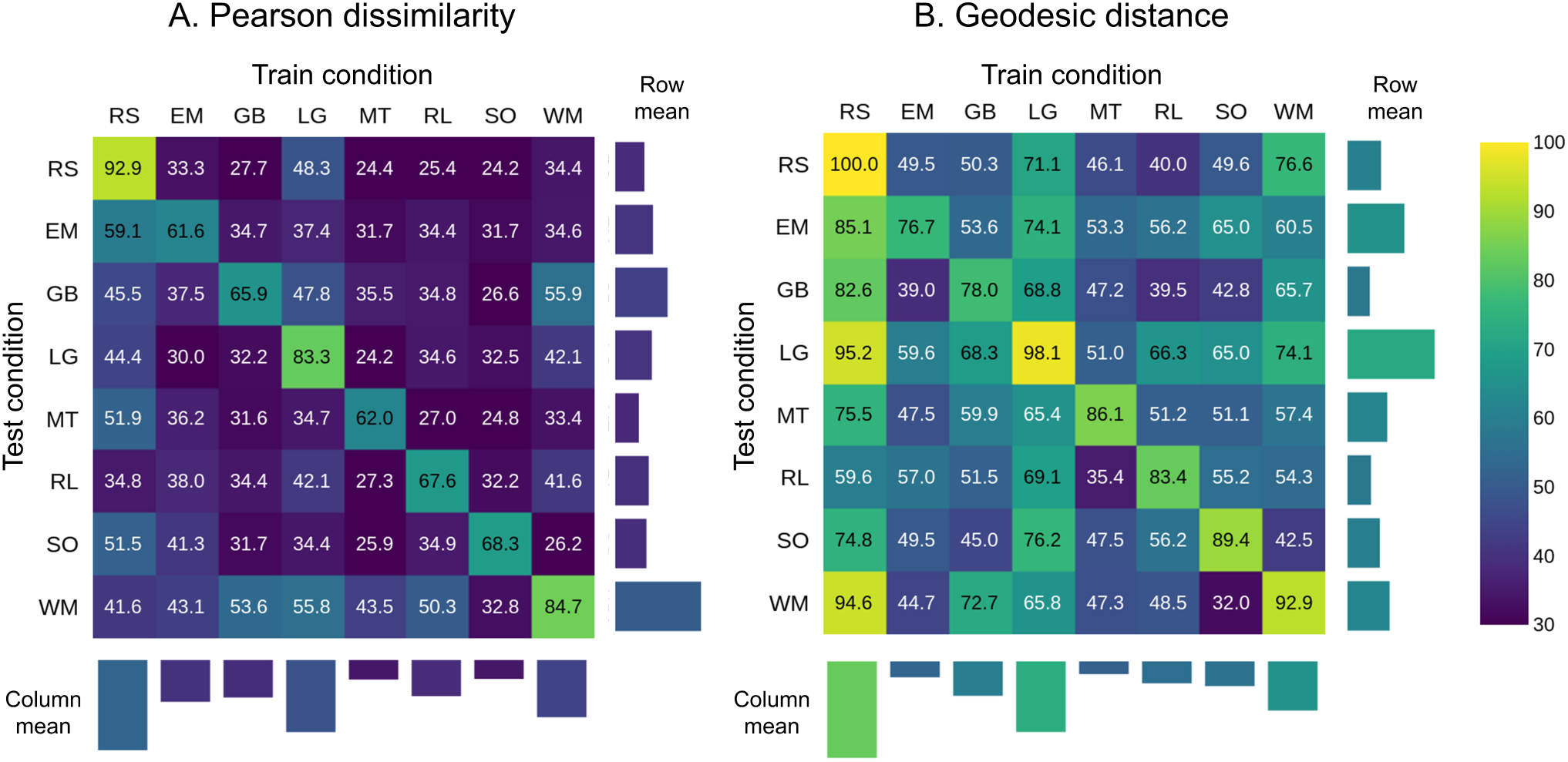
Participant identification accuracy when the training and testing data were based on different conditions. The combined network containing the frontoparietal and default mode subnetworks was employed. The mean accuracy for each train and test condition is also indicated. For example, when *resting-state* is used as training data, the column mean is computed as the accuracy across all other conditions (i.e., except *resting-state* itself). The row means are computed in a similar fashion by excluding the diagonal term. Abbreviations: EM, emotion processing; GB, gambling; LG, language; MT, motor; RL, relational processing; RS, resting-state; SO, social cognition; WM, working memory.

Because in this section time course length was not equated across conditions, we note that those with longer lengths aided across-task identification. Accordingly, transfer might particularly benefit from employing training sets with longer data segments. Nevertheless, future research should also evaluate transfer effects when longer data segments are available for a wider range of tasks (for example, ≥ 300 TRs) so as to characterize their transfer potential.

### 3.9. FC geometry of task and resting-state data

As some conditions yielded high identification accuracy when training and testing were based on different conditions, we sought to visualize distance/dissimilarity in a lower dimensional space via multidimensional scaling. Fig. 12A displays the low-dimensional representation of the distances/dissimilarities for a set of randomly chosen participants when *resting-state* was employed for training data and working memory for testing (untrimmed data). Based on the geodesic distance, *resting-state* FC matrices were relatively close together to one another; in contrast, *working memory* FC matrices were further “spread out”. Intriguingly, such geometry allowed for the separation of FCs based on participant identity. To see this, consider the panels in Fig. 12B, which show participant-level distances. In contrast, using Pearson dissimilarity, the geometry did not allow accurate participant identity. In fact, nearly all participants in this illustrative sample were misidentified.

**Fig. 12:**
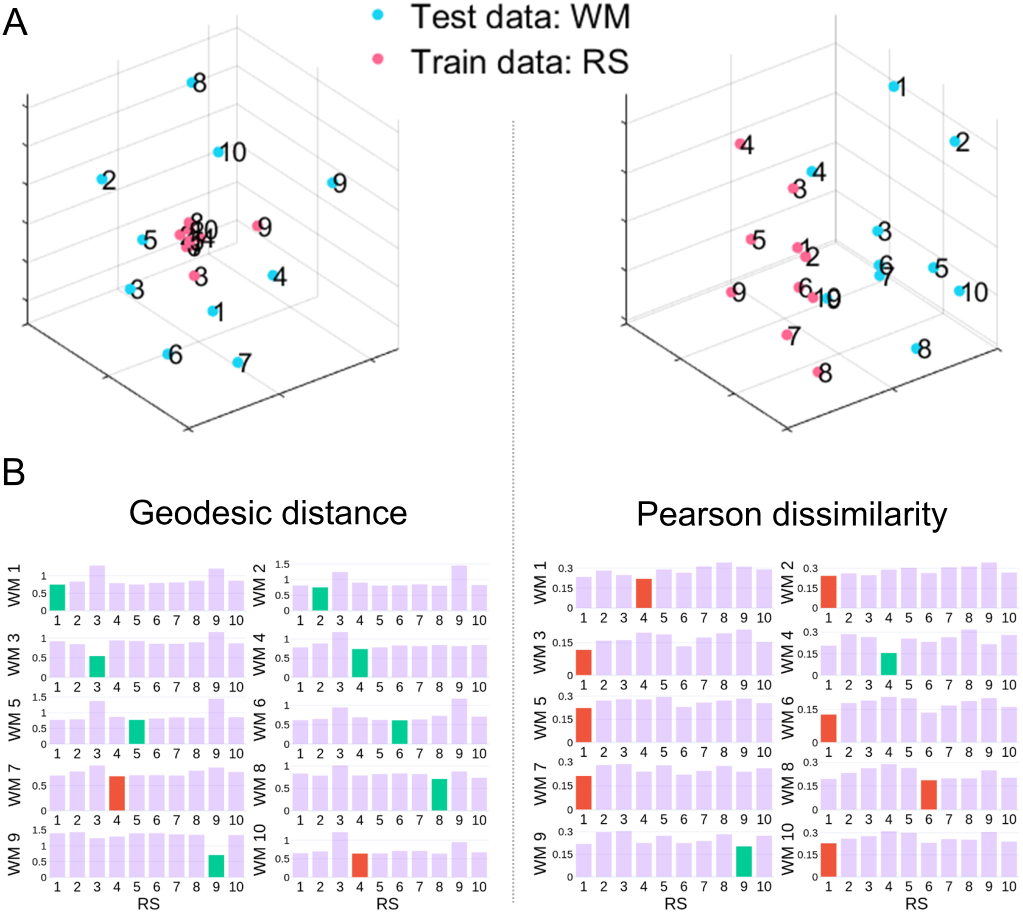
Visualization of task and *resting-state* functional connectivity distances/dissimilarities in a three-dimensional space using multidimensioanl scaling. The numbers indicate participant IDs. (A) Distances/dissimilarities between the functional connectivity matrices of *resting-state* (RS, used for training) and *working memory* (WM, used for testing) for a set of 10 randomly chosen participants. Online figures are available [37]. (B) Participant-level distances/similarities between training and testing data. Correct identification is marked in green and incorrect in red. For example, when using geodesic distance, the best candidate for WM participant 1 (call it WM1) was RS participant 1 (RS1), and the best candidate for WM2 was RS2. However, incorrect classifications were also observed, such as RS4 (not RS7) being closest to WM7. For Pearson dissimilarity most classifications were incorrect, such as RS1 (not RS10) being most similar to WM10. Distances based on the two measures have arbitrary units, and are not comparable across them.

The results in Fig. 12A prompted us to investigate, in an exploratory fashion, distance/dissimilarity between conditions, specifically, *resting-state, motor*, and *language* (Fig. 13). Intriguingly, the geometry of distances was quite different when geodesic distances were used compared to Pearson dissimilarity. These observations suggest that when FC matrices are used for *task classification* (not identification as done here), different algorithms may be more suited for this aim. For example, non-linear radial basis functions might function better for the geodesic case, and linear classifiers for Pearson dissimilarity. Although a fuller investigation of this issue is beyond the scope of the present paper, we believe this is a fruitful direction. Furthermore, the analysis of functional connectivity of mental states should take into account participant-related information since it plays a potentially dominant contribution in the identification of mental states [38].

**Fig. 13:**
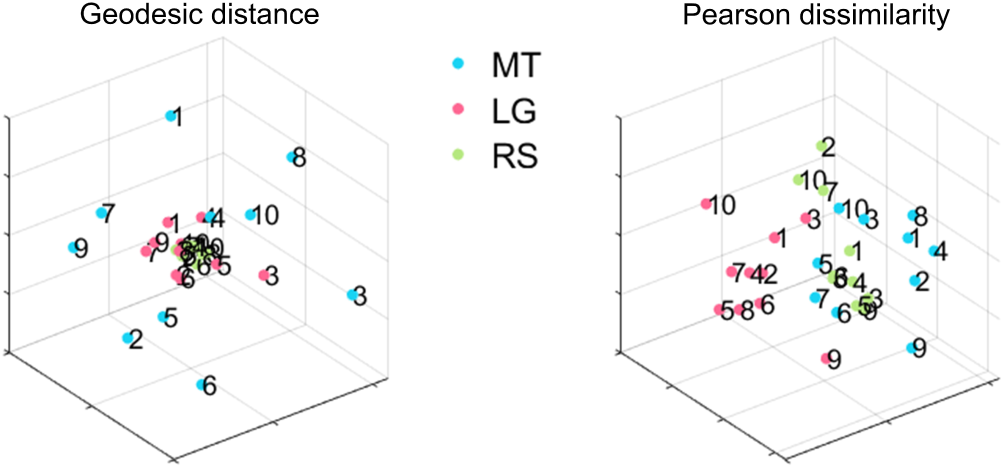
Functional connectivity geometry of *resting-state* and task conditions (online figures are available [37]). Training data for 10 random participants were employed (indicated by the numbers). Distances/dissimilarities in low dimensions were obtained via multidimensional scaling. Note that the geometry in low dimensions differed considerably for geodesic and Pearson, suggesting that condition categorization (not participant identification) should capitalize on such geometry for better performance. Abbreviations: MT, motor; LG, language; RS, resting-state.

## 4. Discussion

In this paper, we investigated participant identification based on FC matrices from fMRI data by employing geometry-aware methods. Correlation matrices are objects that lie on non-linear surfaces, and thereby benefit from non-Euclidean distance measures. Indeed, we show that using the geodesic distance improved participant identification, at times by as much as 20%. Further, low-dimensional visualization based on geodesic distance contributes to understanding how FC geometry affects identification.

### 4.1. Factors influencing participant identification

Scan duration determines the amount of data used to estimate FC matrices, and played a key role in identification accuracy. For *resting-state* data, accuracy improved with time course length and was close to 95% when the entire data were employed (1200 TRs), but fell to under 50% when trimmed to under 150 TRs. The steep drop is possibly due to the underlying dynamics of *resting-state* data [2], and reveals that longer data segments are required to more robustly identify functional connectivity patterns that are unique to individuals. Notably, using the geodesic distance resulted in higher accuracy than Pearson dissimilarity even when, say, only a fourth of the data were employed for FC estimation. Thus, a more suitable geometry is particularly appealing when data-limited scenarios are envisioned.

When time course length was trimmed to the same duration, identification accuracy still varied across scanning conditions. The *resting-state* condition resulted in the lowest accuracy. With the data trimmed to the minimum amount of data, the *language* task exhibited over 80% accuracy. Accuracy of all task conditions exceeded 50%, with four of them exceeding 60%. Thus, even with rather limited amounts of data identifying the participant was considerably better than the chance level of 1%. In addition, we observed considerable variability is performance across conditions, consistent with previous literature suggesting that brain states can be manipulated to emphasize individual differences in FC [14].

Thus far, we have discussed findings based on whole-cortex FC matrices (300 ROIs were employed). We reasoned that particular subsets of regions potentially might be more informative than others. To evaluate this possibility identification was applied to *resting-state* and task conditions separately for each individual subnetwork of the Yeo parcellation ([39]). The FC matrices employed were therefore relatively small (the number of ROIs ranged from 20-68). Four subnetworks (`vision, dorsal-attention, frontoparietal, default`) stood out as consistently exhibiting the highest levels of performance. The average accuracy across conditions approach 70% for the four networks. Intriguingly, accuracy for the *language* task based on the `frontoparietal` and `default` subnetworks exceeded that observed with the whole cortex. Whereas subnetwork size might contribute to its ability to identify participants, it is clearly not the driving factor. For example, the `dorsal-attention` and the `ventral-attention` networks had the same number of ROIs, but the former outperformed the latter consistently (on average by over 30%).

To further explore subnetwork contributions we also combined the two that displayed the highest individual accuracy (`frontoparietal` and `default`) into a single network. Remarkably, the combined network always numerically outperformed the individual subnetworks, and indeed the entire cortex. When additional subnetworks were combined, accuracy initially increased but then decreased. Accuracy peaked at two or three subnetworks, with whole-cortex FCs having the worst performance across conditions. Accuracy also varied across combinations of subnetworks, with the minimum value less than half that of the maximum. These results are related to the non-uniformity of within-subject test-retest reliability of connectivity profiles, and might inform how individual differences are associated with heritability and cognitive ability [11]. Thus, future work on individual differences using connectomes should not only consider tasks but also choose appropriate measures and subnetworks that emphasize these differences.

Although it was beyond the scope of the present study, it would be valuable to investigate in future studies factors contributing to the performance of individual subnetworks, and their combinations. For example, subnetworks may contribute highly to identification because their individual-specific functional connectivity information capitalizes on the contributions of these subnetworks to task performance. Alternatively, but not mutually exclusively, subnetworks that do not participate as much during a task may contain diagnostic information with respect to participant identity.

To what extent does participant identification transfer between experimental conditions? We found that training with one condition and testing with another produced good levels of identification accuracy. Certain combinations that on the surface were not obvious produced particularly impressive results; for example, training with *gambling* and testing with *working-memory*, or training with *working-memory* and testing with *language*. Training with *motor* produced the least transfer to other tasks, perhaps due to the low-level specificity of this task. Notably, training with *resting-state* produced very high transfer, such that testing with each task attained accuracy over 75% (with the exception of *relational processing*), and in some instances over 90%. The choice of measure was particularly important for transfer of identifiability and accuracy, with *working-memory* attaining nearly 95% using geodesic distance but less than 42% using Pearson dissimilarity.

### 4.2. Low-dimensional distance visualizations

Relationships between high-dimensional FC matrices (300 × 300) were visualized in three Euclidean-space dimensions using multidimensional scaling. Both the Pearson dissimilarity measure and geodesic distance were used. Note that computing geodesic distances takes into account the non-linear geometry of correlation matrices. Once their distances are computed, and the space nonlinearity taken into account, they can be illustrated in Euclidean space (naturally, some distortion ensues due to dimensionality reduction).

In our explorations, low-dimensional visualizations reflected identification accuracy on the full data, and thus preserved important distance information. In particular, the higher identification accuracy using the geodesic distance resulted in relatively low within- and high between-participant distances. Visualization of FC from task data revealed insights into the geometry of task correlation matrices in relation to *resting-state*. Identification accuracy is related to the ratio of within- to between-participant distances. Surprisingly, with geodesic distances, tasks associated with higher identification accuracy exhibited smaller between-participant distances. Still, the more favorable ratio of within- to between-participant distances led to favorable identification accuracy. Thus, the underlying geometry of functional connectivity may provide further insights into our finding that high identification accuracy was attained when training and testing were based on different scanning conditions.

In the visualizations based on geodesic distance, distances between task FCs did not appear to form convex sets (if *A* and *B* are two points in a convex set, any point on the line joining them also belongs to the set), and were instead in clustered arrangements. Of note, previous work performing clustering of FCs [2, 18] have used *k*-means which are not well suited to finding non-convex clusters [13]. Instead, methods such as spectral clustering [27] and non-linear support-vector kernels [9] are capable of capturing very general structures, and are potentially more suitable for classifying functional connectivity.

Pearson correlation is a common approach to compare FC matrices. The present study demonstrates that non-linear measures are better suited to characterize functional connectivity relationships. The low-dimensional visualization briefly explored here hints at the different geometries associated with the geodesic non-linear metric and the Pearson approach. Surprisingly, we noted in our investigations that simple visual inspection of the correlation matrices as commonly done in the field to highlight similarities between conditions can also be problematic, and in fact can lead to unintuitive scenarios (Fig 14).

**Fig. 14:**
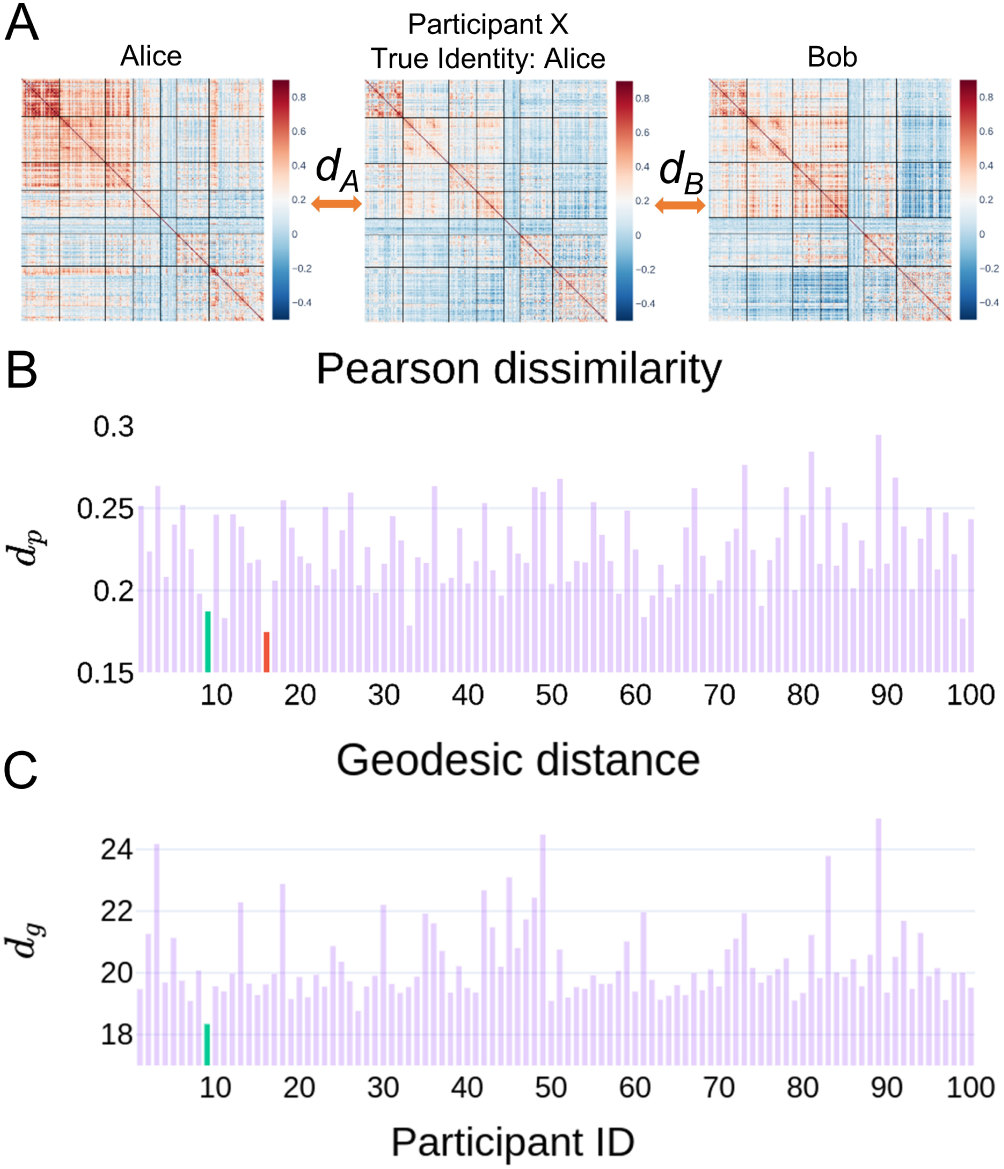
Visual comparison of functional connectivity (FC) matrices can be unintuitive. (A) Example FCs from *resting-state* data where the geodesic distance correctly labeled the test participant but Pearson dissimilarity did not. Pearson dissimilarities and geodesic distances between the test-FC and each of the FCs in the training data are shown in (B) and (C). The green bar indicates the distance between the test-FC to the correct training set FC; the red bar indicates an incorrectly labeled training set FC. For the geodesic distance, the labeled participant had indeed the smallest value; not so in the case of Pearson dissimilarity. This example also questions the common practice of informally evaluating functional connectivity similarity via simple visual inspection. At the very least, it is not immediate that participant *X* is more similar to Alice than Bob.

### 4.3. Conclusions

Time series correlation matrices capture important aspects of brain functional organization. Here, we propose the use of a geodesic distance metric that reflects the underlying non-Euclidean geometry of functional connectivity matrices. We compared identification performance (also called “fingerprinting”; that is, assigning a participant label to novel functional connectivity data) obtained with standard Pearson correlation and the proposed geodesic distance. The latter not only improved identification accuracy but also provided insights into the geometry of task and resting-state conditions. Importantly, the approach advocated here is general and can be utilized to study the clustering of brain states, how tasks potentially reconfigure brain networks, and to characterize intersubject correlations. Code and html figures are available at https://github.com/makto-toruk/FC_geodesic.

## Acknowledgements

M.V. was supported by a fellowship by the Brain and Behavior Initiative, University of Maryland, College Park. L.P. is supported by the National Institute of Mental Health (R01 MH071589 and R01 MH112517). Data were provided by the Human Connectome Project, WU-Minn Consortium (Principal Investigators: David Van Essen and Kamil Ugurbil; 1U54MH091657) funded by the 16 NIH Institutes and Centers that support the NIH Blueprint for Neuroscience Research; and by the McDonnell Center for Systems Neuroscience at Washington University. The authors would like to thank Jingwei Li, Ru Kong, and Thomas Yeo for providing resting-state data that included global signal regression in the preprocessing pipeline.

---

**Fig. S1:**
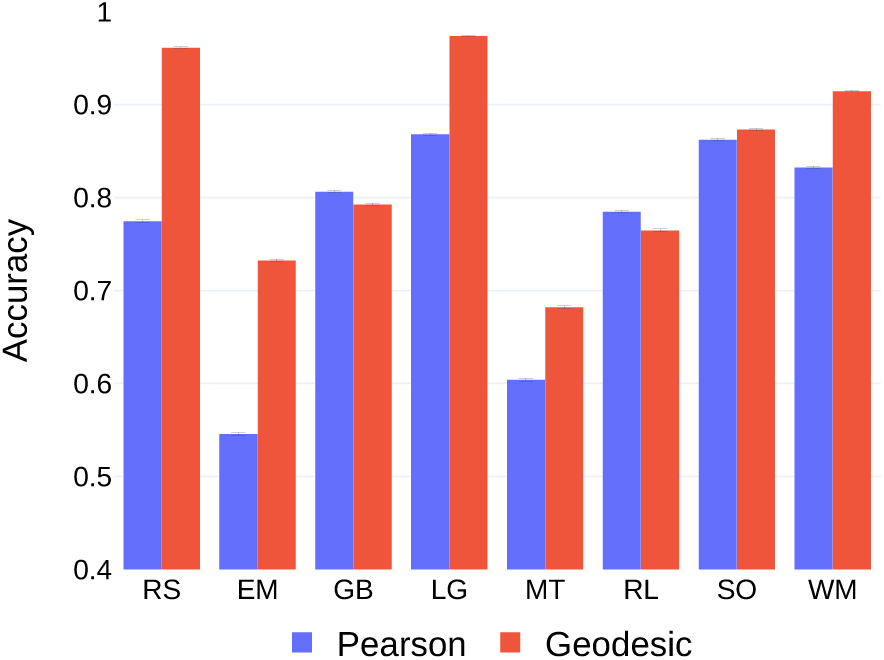
Participant identification for the eight conditions using the geodesic distance and Pearson dissimilarity. Training and testing data were from the same condition. Fixation period or “mini resting periods” were not trimmed from runs of task data (as done in Fig. 3). Abbreviations: EM, emotion processing; GB, gambling; LG, language; MT, motor; RL, relational processing; RS, resting-state; SO, social cognition; WM, working memory.

## Supplemental material

### S1. Identification accuracy when runs were not trimmed

In the main body of the text, to ensure that only task-related segments of a run were retained, “mini resting periods” in the form of fixation periods were removed (see Section 2.1). We repeated our analysis without trimming runs of task data using whole-cortex FCs. Identification accuracy for each condition is shown in Fig. S1. Accuracy obtained using the geodesic distance exceeded that of Pearson dissimilarity for all conditions except the *gambling* and *relational* tasks (*p* = 1 for *gambling* and *relational* tasks, *p* < 10^−6^ for all tasks; reference *α* = 0.05/8 = 0.00625 given 8 conditions; Fig. S2). The mean improvement using geodesic distance was around 8% (as high as 18% on *resting-state* data).

### S2. Effect of global signal regression on identification

We repeated our analysis by including global signal regression (GSR) in the preprocessing pipeline for resting-state data [20, 23]. The use of GSR is still debated [26] and can potentially spread underlying group differences to regions that may never have had any [31]. We limit our analysis in this section to *resting-state* data; we did not include GSR in the preprocessing pipeline for results in the main text. In the data employed (see Acknowledgements), 8 subjects’ data were removed because they did not not pass quality control check [23]. Thus, the results reported this section were based on *N* = 92 participants. We performed participant identification using whole-cortex FCs. Regardless of the inclusion of GSR in preprocessing, identification accuracy improved using geodesic distance compared to Pearson dissimilarity (Fig. S3A). However, using GSR improved accuracy for both measures. When segments of smaller lengths were extracted from *resting-state* data, accuracy improved using geodesic distance for all segment lengths (Fig. S3B). When GSR was used, accuracy using geodesic distance was close to 95% with only 200 time points (compared to 70% without GSR; Fig. S3C).

### S3. Effect of number of ROIs in the parcellation on identification

To study the effect of the parcellation scheme on participant identification accuracy using the two measures, we employed various parcellations with ROIs ranging from a 100 to 400. In general, mean participant identification accuracy increased with increase in ROIs indicating that finer resolution or detail in the FC revealed more uniqueness. Mean accuracy using the geodesic distance was consistently higher than the mean accuracy using Pearson dissimilarity. For several conditions (*resting-state, language, motor*), accuracy using geodesic distance on FCs obtained with 100 ROIs was greater than accuracy obtained using Pearson dissimilarity with 400 ROIs (Fig. S5).

### S4. Computing geodesic distances for matrices without full rank

Computing the geodesic distance between two FC matrices *Q*_1_ and *Q*_2_ (Equation 3) requires *Q*_1_ to be invertible, or equivalently, all the eigenvalues of *Q*_1_ must be strictly greater than zero. When FC matrices are based on *n* ROIs and *n* is larger than number of frames in the run, the *rank* of the resulting FC matrix is not full (i.e., < *n*), and some of its some eigen-values are equal to 0. In practice, when the number of ROIs *n* < (0.9 × number of frames), we applied the procedure below to ensure full rankness.

To handle such cases, we adopted a simple approach here: we added the identity matrix *I* to both *Q*_1_ and *Q*_2_, causing the eigenvalues of the correlation matrices of interest to be increased by 1. Because all eigenvalues are then greater than 0, the matrices are invertible. In such cases, the geodesic distance, *d*_*G*_(*Q*_1_+*I, Q*_2_+*I*), serves as a proxy for the geodesic distance between the two matrices. Note that the scenario of low-rank FC matrices arises only for whole-cortex analysis, as for the subnetwork analyses, the number of ROIs in question was always greater than the number of frames in the run.

For reference, the procedure above was employed in the following cases: whole-cortex results for all tasks; whole-cortex *resting-state* results with lengths less than 400 TRs; and whole-cortex results involving trimmed data.

---

**Fig. S2:**
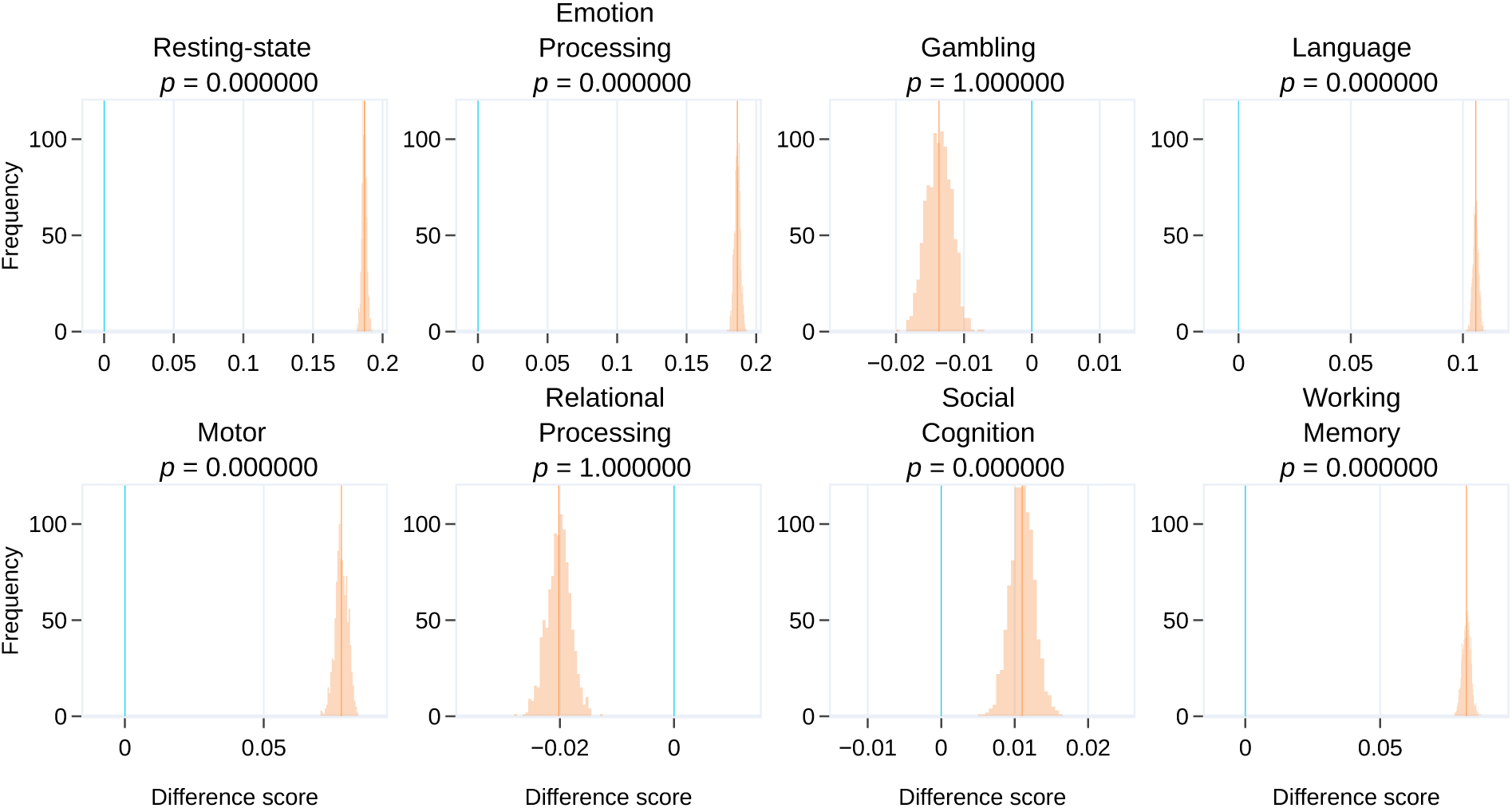
Whole-cortex FCs without trimming runs: Comparison of identification accuracy based on geodesic distance and Pearson dissimilarity for each condition. Identification was based on whole-cortex FCs. Runs were not trimmed as in the main body of the work (see Section 2.1). For each condition, the distributions shown in orange represent the difference between the mean participant identification accuracy using the geodesic distance and Pearson dissimilarity across the outer bootstrap iterations (see Section 2.7). The orange line indicates the mean of the difference distribution and the blue line indicates zero difference.

**Fig. S3:**
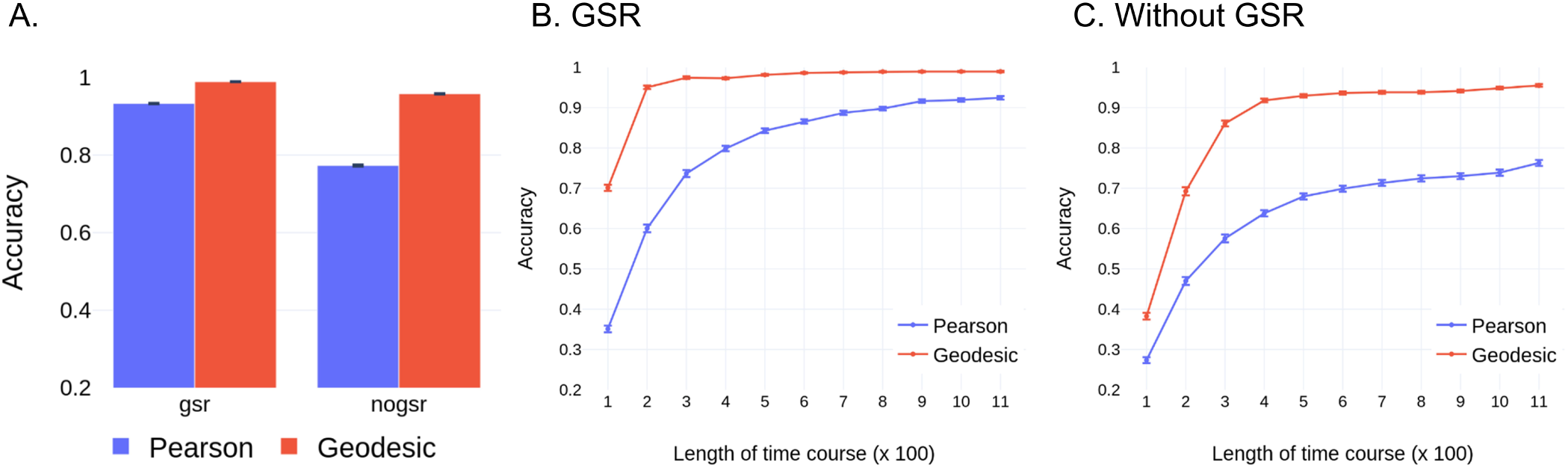
A. Participant identification accuracy with global mean regression (GSR) included in the preprocessing pipeline (gsr) or not (nogsr). Accuracy using geodesic distance exceeded Pearson dissimilarity for both preprocessing methods. Participant identification accuracy as a function of segment length for *resting-state* data with GSR (in B) and without GSR (in C). In both cases, accuracy using geodesic distance exceeded Pearson dissimilarity at each segment length. Error bars indicate standard error of the mean across bootstrap iterations.

**Fig. S4:**
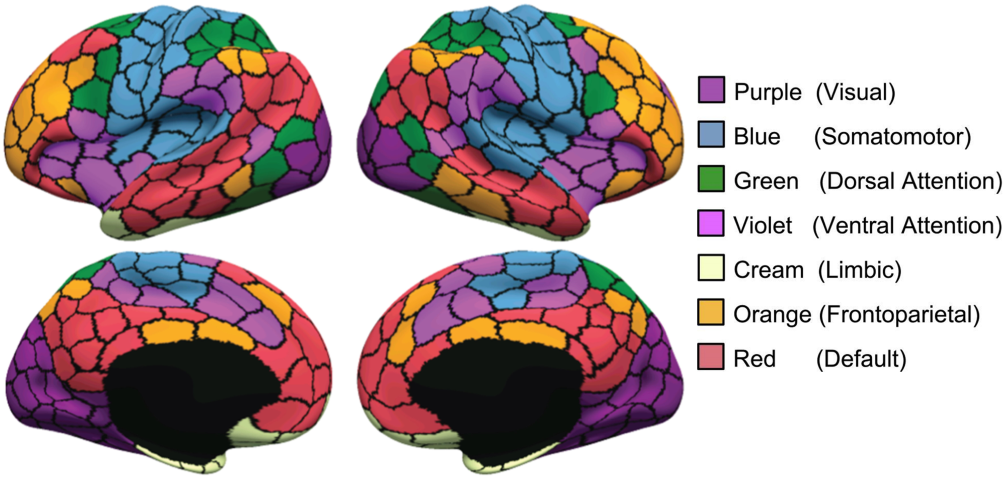
Parcellation of the cortex into 300 ROIs as provided by [32]. ROIs were grouped into the 7 networks described in [39].

**Fig. S5:**
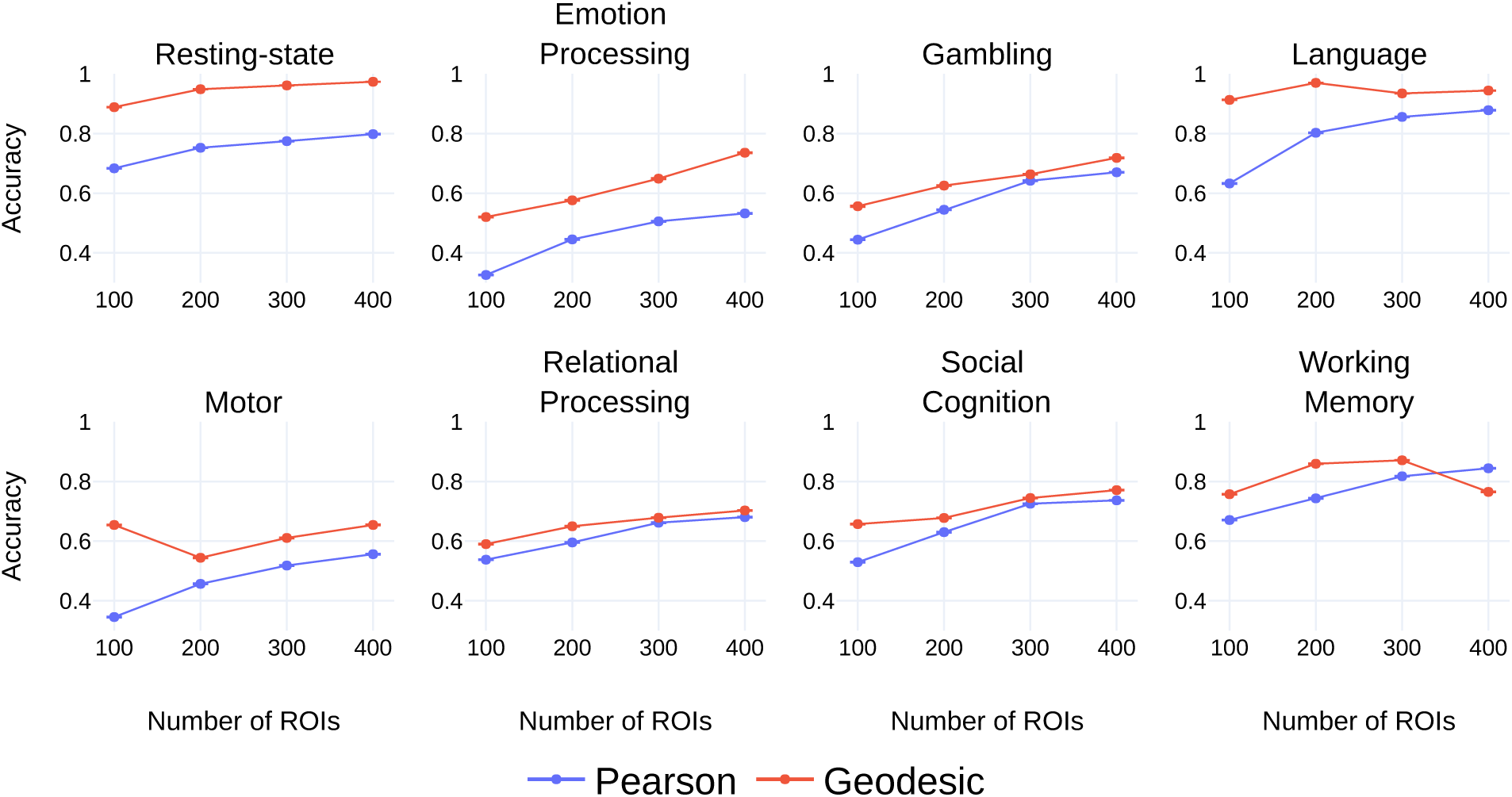
Participant identification accuracy as a function of the number of ROIs. Here, training and testing data are from the same condition. Error bars indicate standard error of the mean across the bootstrap iterations.

**Fig. S6:**
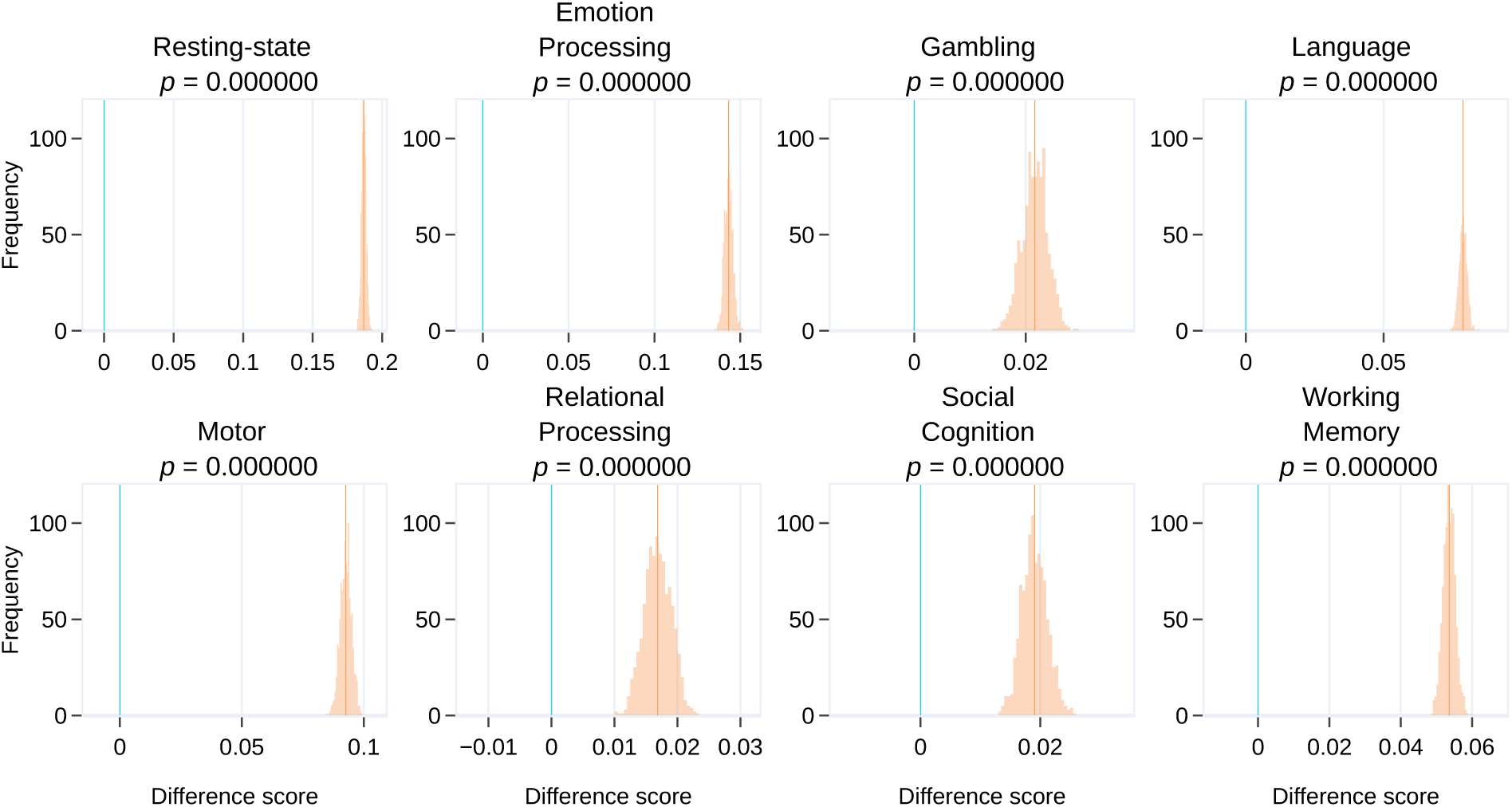
Whole-cortex FCs with full time course lengths: Comparison of identification accuracy based on geodesic distance and Pearson dissimilarity for each condition. Identification was based on whole-cortex FCs. Here, full time course lengths were used (see Section 2.1). For each condition, the distributions shown in orange represent the difference between the mean participant identification accuracy using the geodesic distance and Pearson dissimilarity across the outer bootstrap iterations (see Section 2.7). The orange line indicates the mean of the difference distribution and the blue line indicates zero difference.

**Fig. S7:**
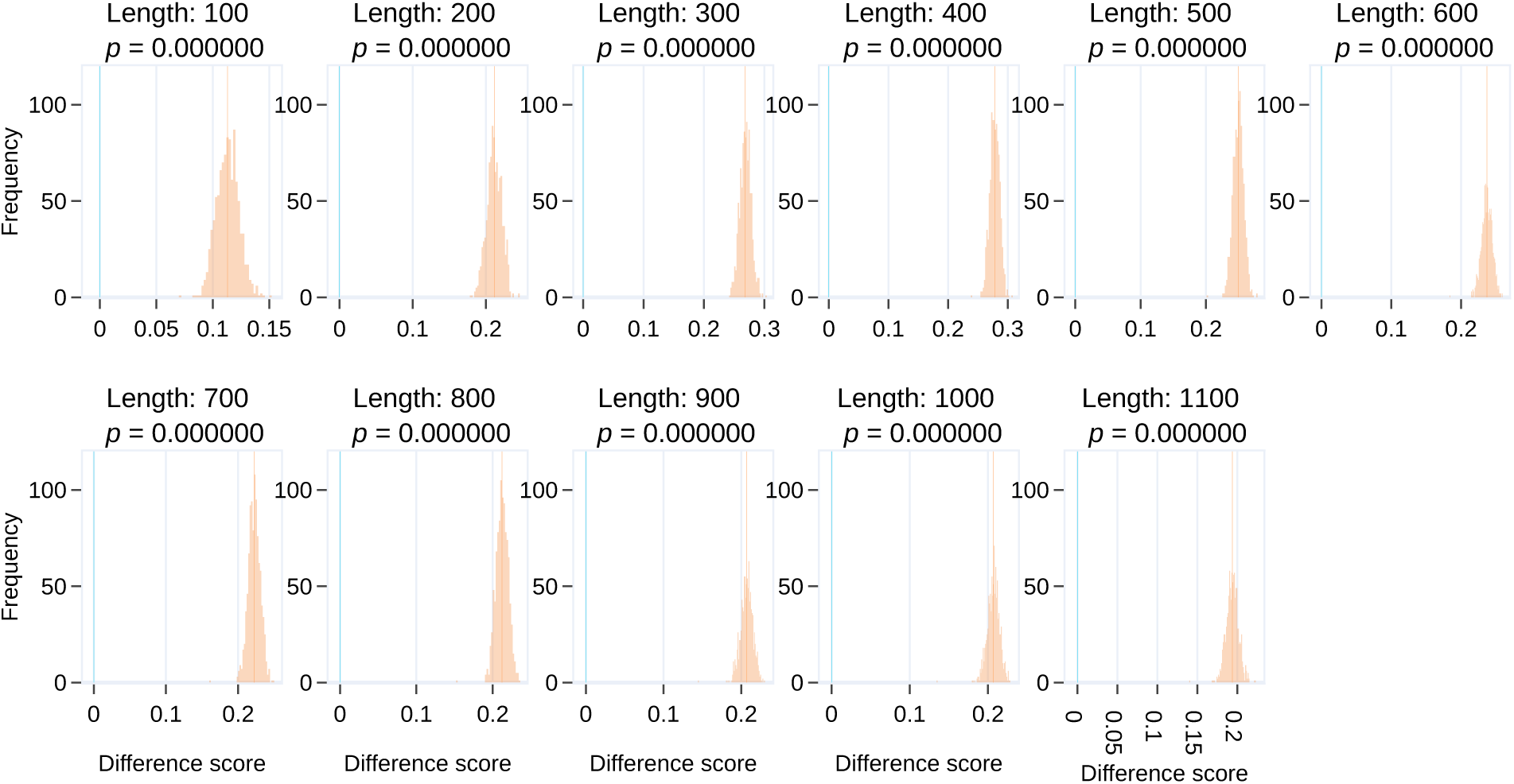
Identification accuracy and time course length: Comparison of identification accuracy based on geodesic distance and Pearson dissimilarity for various time course lengths. Since *resting-state* data had the highest time course length, smaller segments of various lengths were extracted (see Section 2.7.1). Identification was based on whole-cortex FCs. For each segment length, the distributions shown in orange represent the difference between the mean participant identification accuracy using the geodesic distance and Pearson dissimilarity across the outer bootstrap iterations (see Section 2.7). The orange line indicates the mean of the difference distribution and the blue line indicates zero difference.

**Fig. S8:**
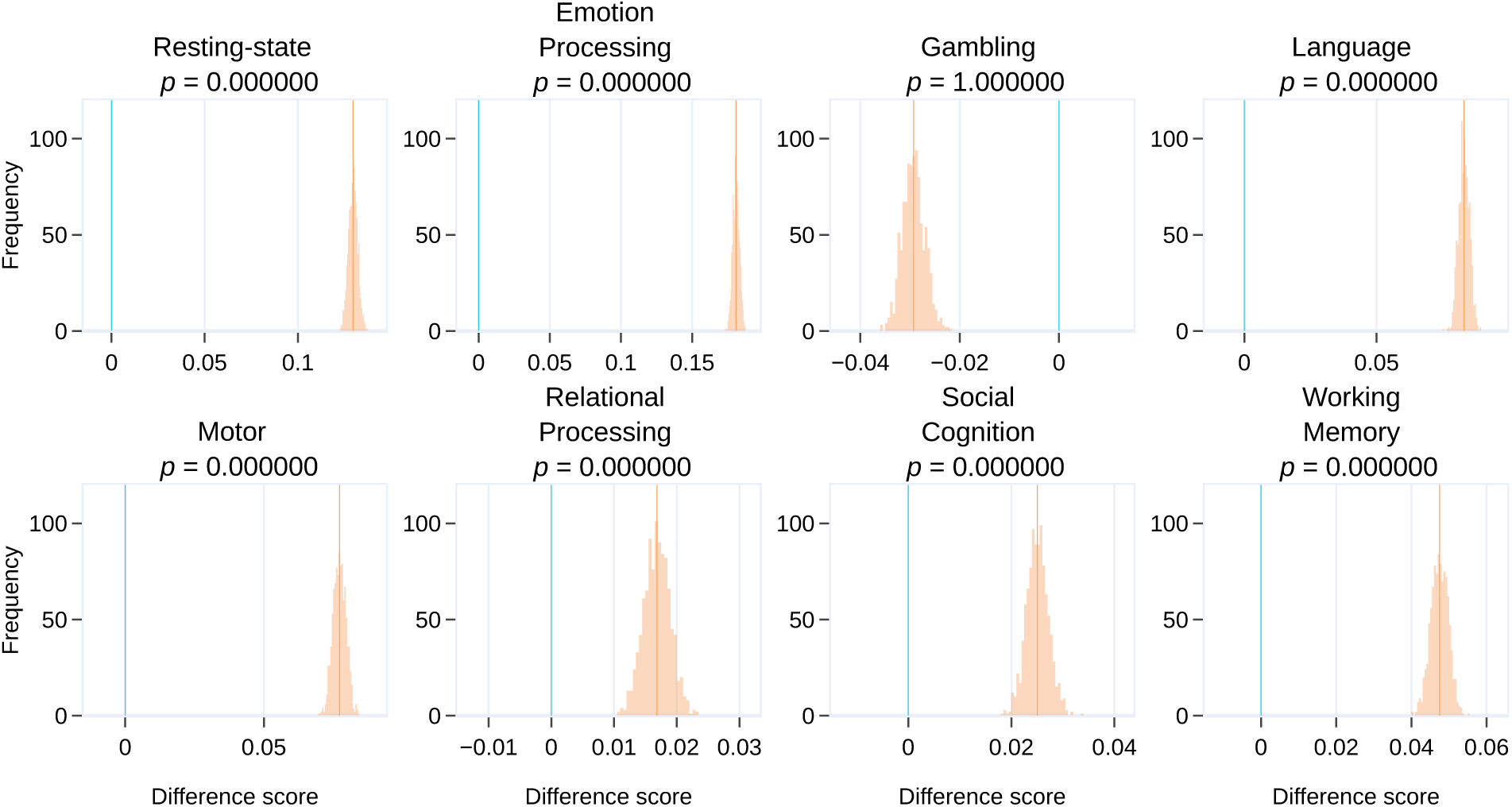
Whole-cortex FCs with trimmed time course lengths: Comparison of identification accuracy based on geodesic distance and Pearson dissimilarity for each condition. Identification was based on whole-cortex FCs. Data for each condition were trimmed such that they all had the same time course length (of 138; see Section 3.5). For each condition, the distributions shown in orange represent the difference between the mean participant identification accuracy using the geodesic distance and Pearson dissimilarity across the outer bootstrap iterations (see Section 2.7). The orange line indicates the mean of the difference distribution and the blue line indicates zero difference.

**Fig. S9:**
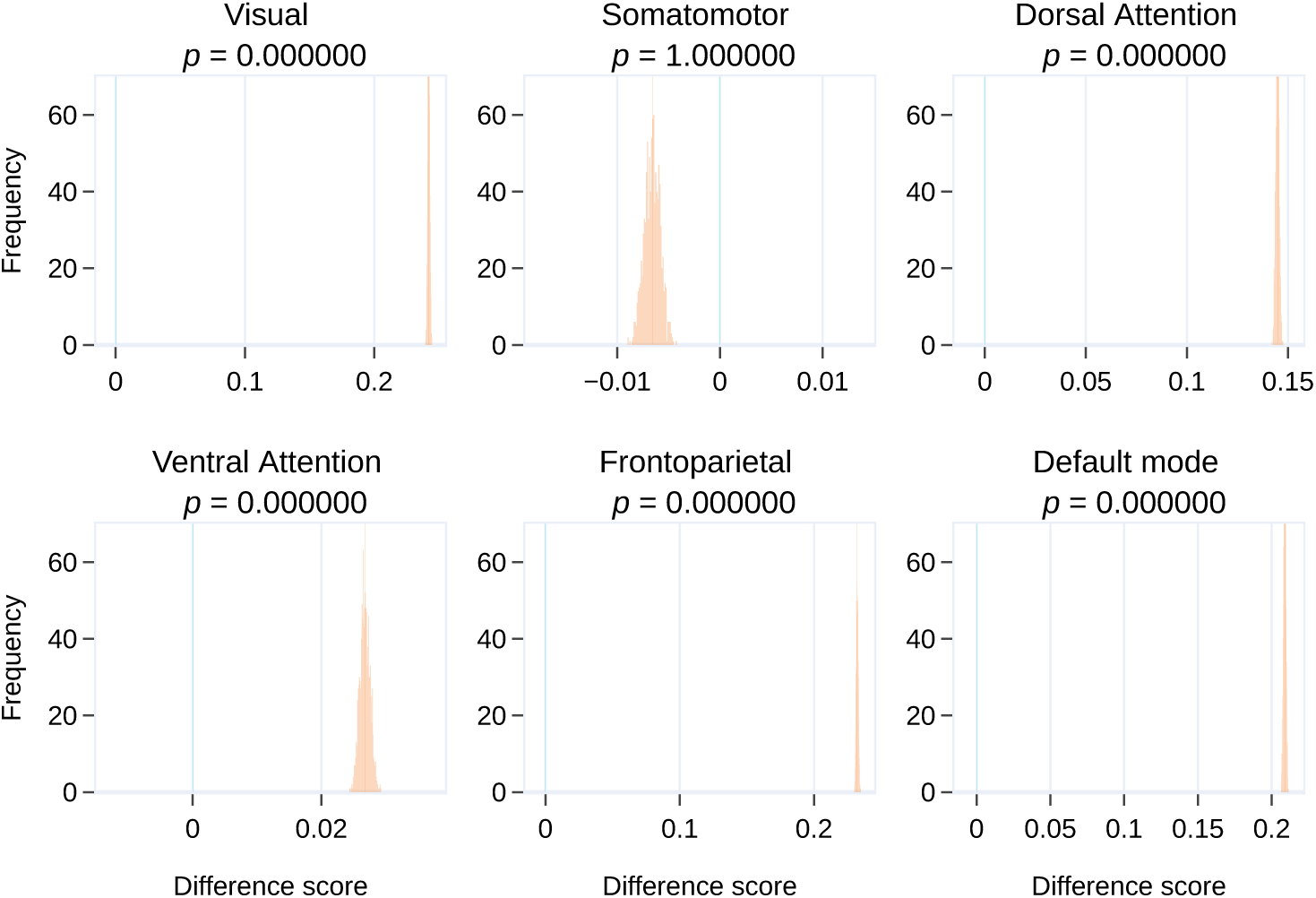
Subnetwork FCs with trimmed time course lengths: Comparison of identification accuracy based on geodesic distance and Pearson dissimilarity for each subnetwork. Identification was based on subnetwork FCs. Data for each condition were trimmed such that they had the same time course length (of 138; see Section 3.5). For each subnetwork, difference scores were averaged across all conditions. The distributions shown in orange represent the difference between the mean participant identification accuracy using the geodesic distance and Pearson dissimilarity across the outer bootstrap iterations (see Section 2.7). The orange line indicates the mean of the difference distribution and the blue line indicates zero difference.

**Fig. S10:**
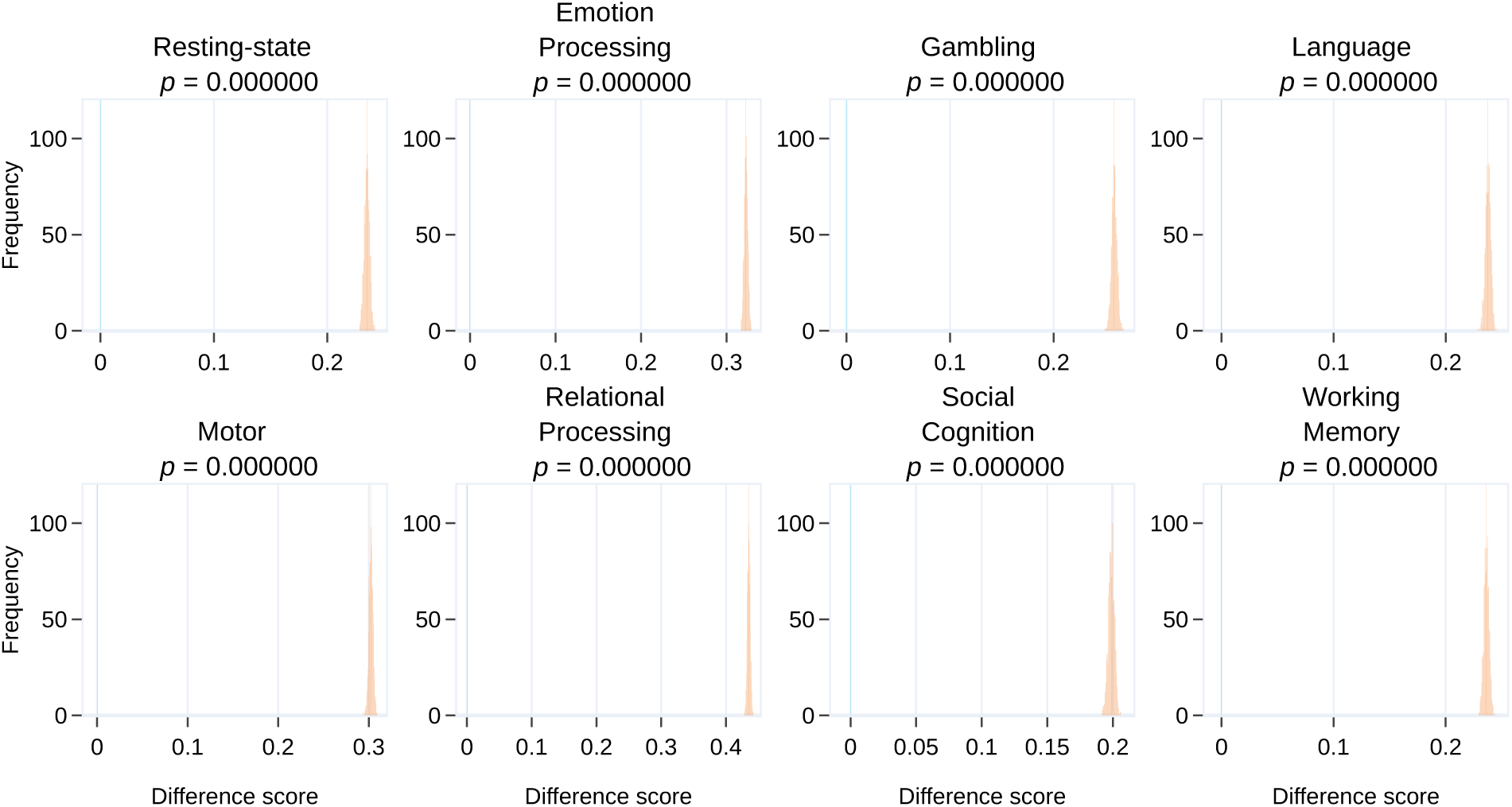
Subnetworks of the same size: Comparison of identification accuracy using `dorsal attention` and `ventral attention` subnetwork FCs for each condition. The geodesic distance measure was used for identification. The two subnetworks were of identical size for the 300 ROIs parcellation (see Table 2. Data for each condition were trimmed such that they all had the same time course length (of 138; see Section 3.5). For each condition, the distributions shown in orange represent the difference between the mean participant identification accuracy based on the two subnetworks across the outer bootstrap iterations (see Section 2.7). The orange line indicates the mean of the difference distribution and the blue line indicates zero difference.

**Fig. S11:**
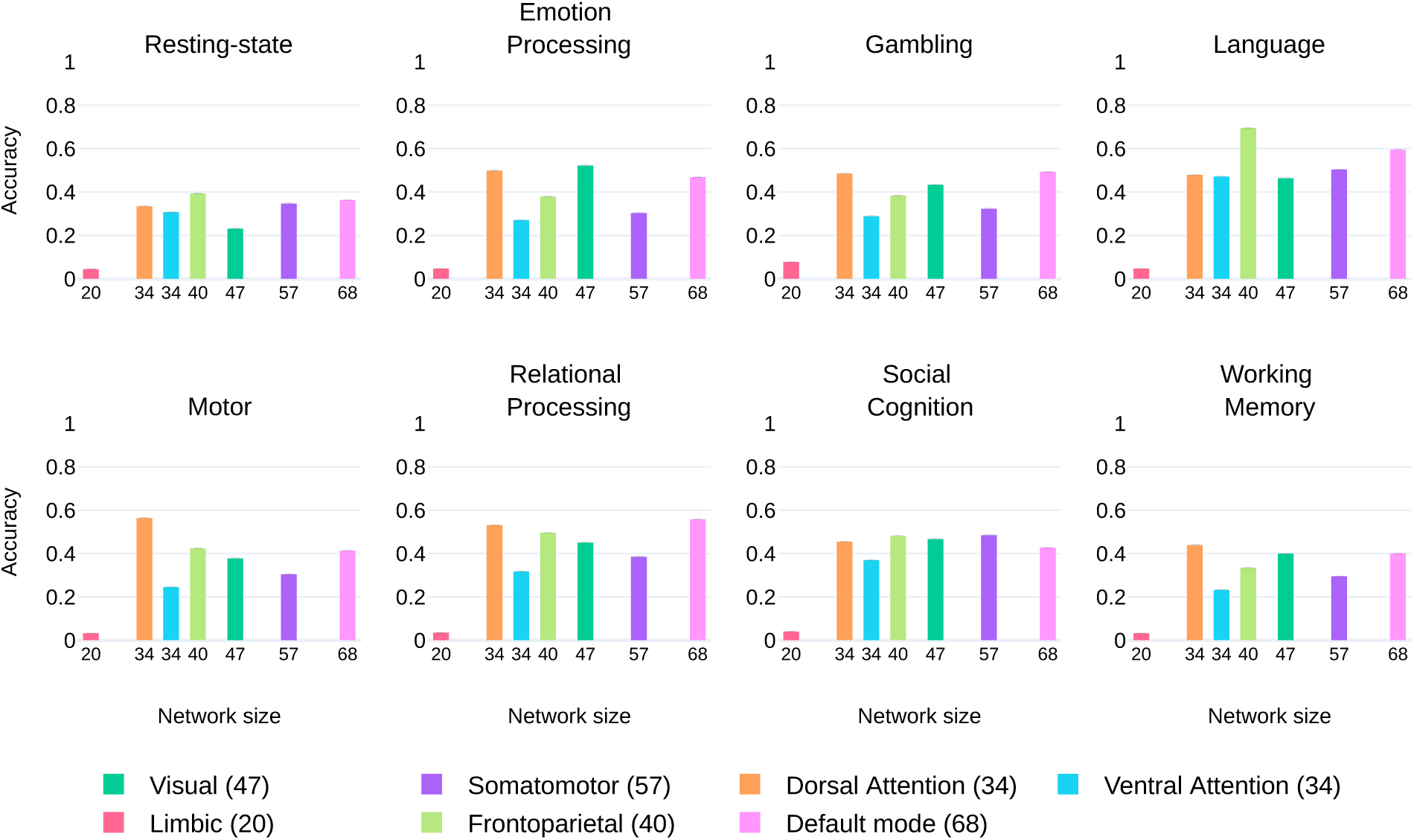
Participant identification accuracy plotted against subnetwork size for each condition (Pearson dissimilarity). The size of the subnetwork (the number of ROIs) is also indicated in the inset. The error bars represent standard error of the mean across bootstrap iterations.

**Fig. S12:**
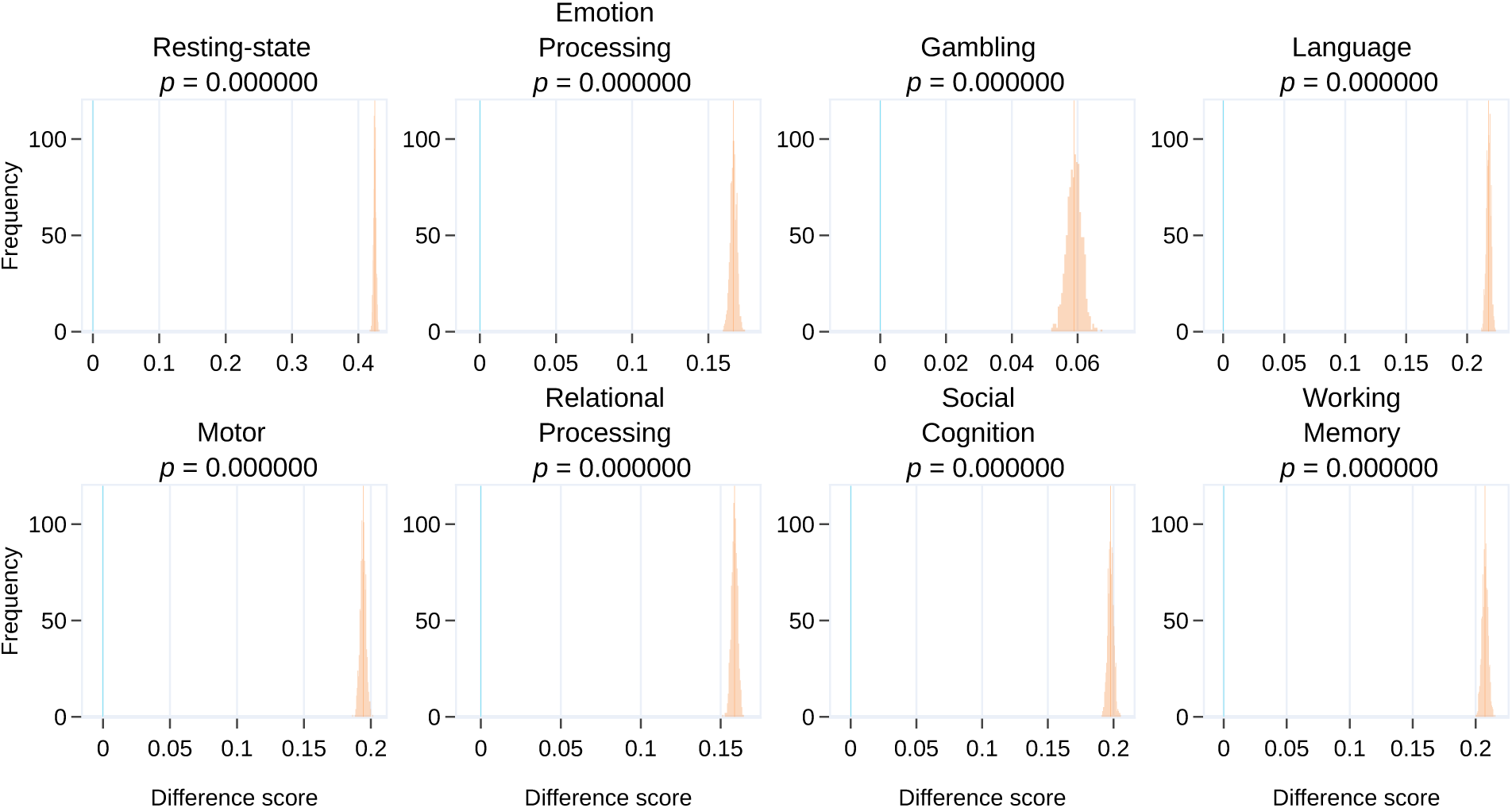
Combined subnetwork FCs with trimmed time course lengths: Comparison of identification accuracy based on geodesic distance and Pearson dissimilarity for each condition. Identification was based on combined subnetwork FCs (see Section 3.7). Data for each condition were trimmed such that they all had the same time course length (of 138; see Section 3.5). For each condition, the distributions shown in orange represent the difference between the mean participant identification accuracy using the geodesic distance and Pearson dissimilarity across the outer bootstrap iterations (see Section 2.7). The orange line indicates the mean of the difference distribution and the blue line indicates zero difference.

**Fig. S13:**
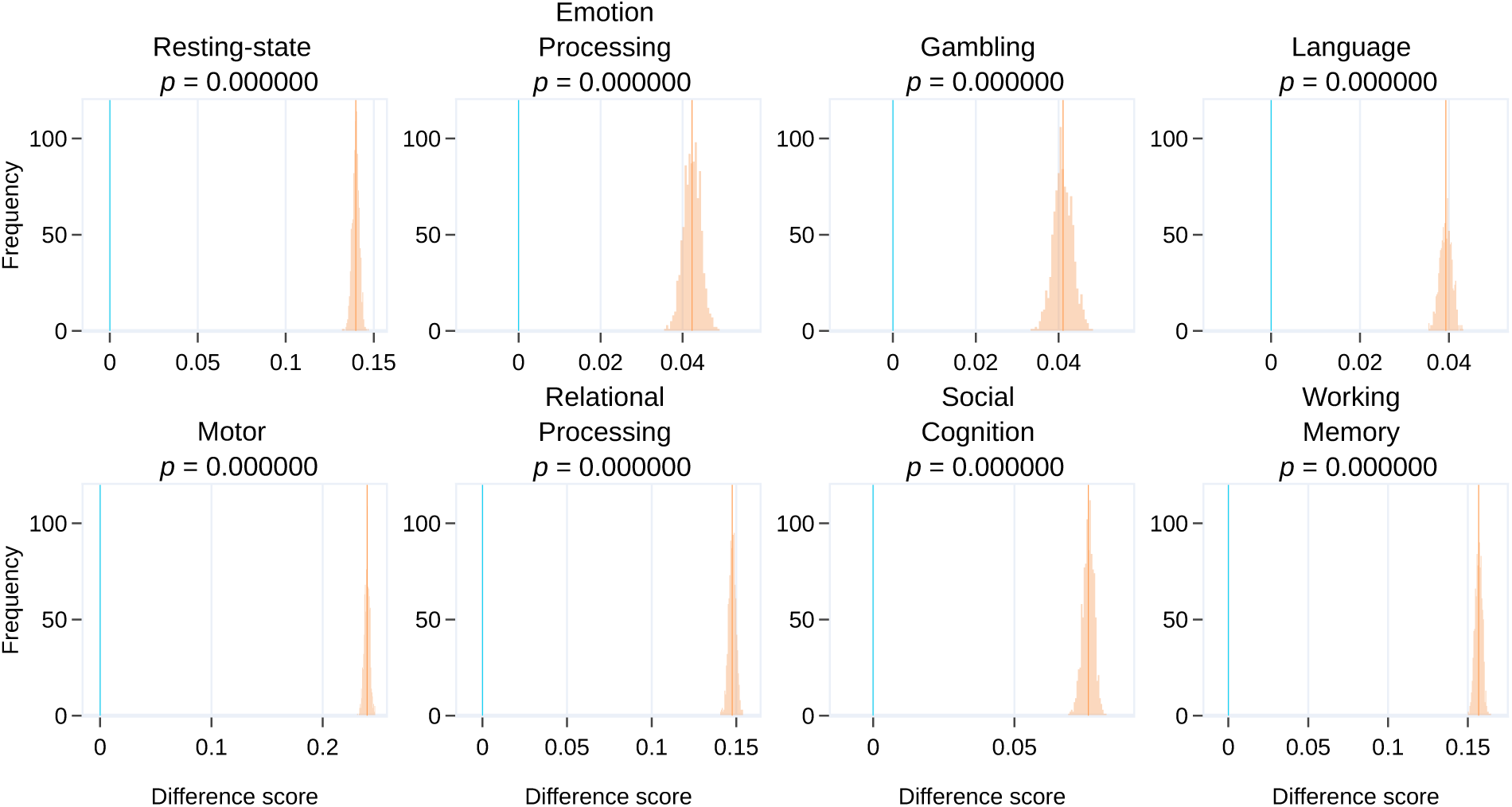
Combined subnetwork vs frontoparietal: Comparison of identification accuracy based on combined subnetwork FCs (see Section 3.7) and `frontoparietal` subnetwork FCs (part of the combined subnetwork) for each condition. The geodesic distance measure was used for identification. Data for each condition were trimmed such that they all had the same time course length (of 138; see Section 3.5). For each condition, the distributions shown in orange represent the difference between the mean participant identification accuracy based on combined subnetwork FCs and `frontoparietal` FCs across the outer bootstrap iterations (see Section 2.7). The orange line indicates the mean of the difference distribution and the blue line indicates zero difference.

**Fig. S14:**
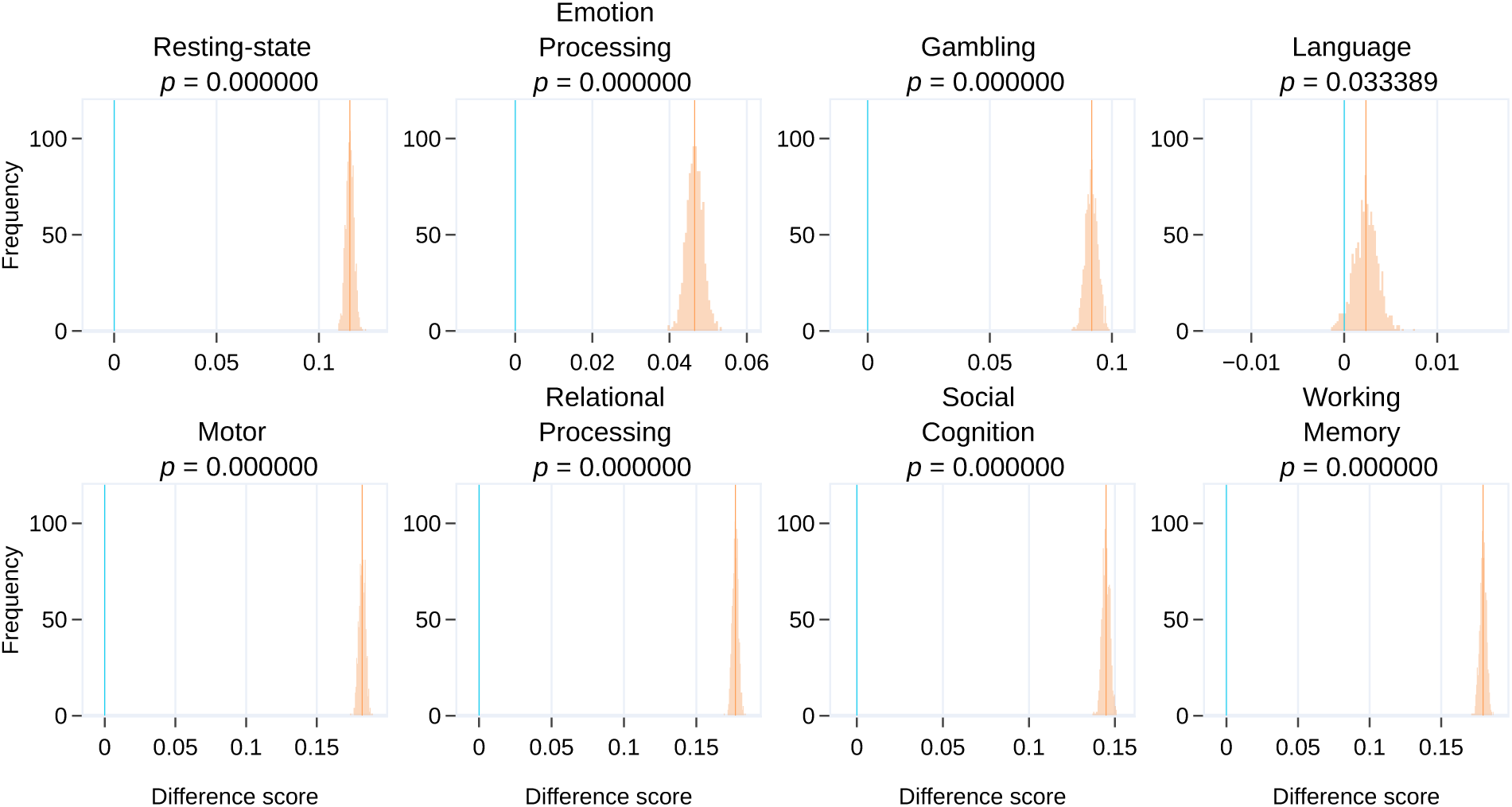
Combined subnetwork vs default mode: Comparison of identification accuracy based on combined subnetwork FCs (see Section 3.7) and default mode subnetwork FCs (part of the combined subnetwork) for each condition. The geodesic distance measure was used for identification. Data for each condition were trimmed such that they all had the same time course length (of 138; see Section 3.5). For each condition, the distributions shown in orange represent the difference between the mean participant identification accuracy based on combined subnetwork FCs and `default mode` FCs across the outer bootstrap iterations (see Section 2.7). The orange line indicates the mean of the difference distribution and the blue line indicates zero difference.

**Fig. S15:**
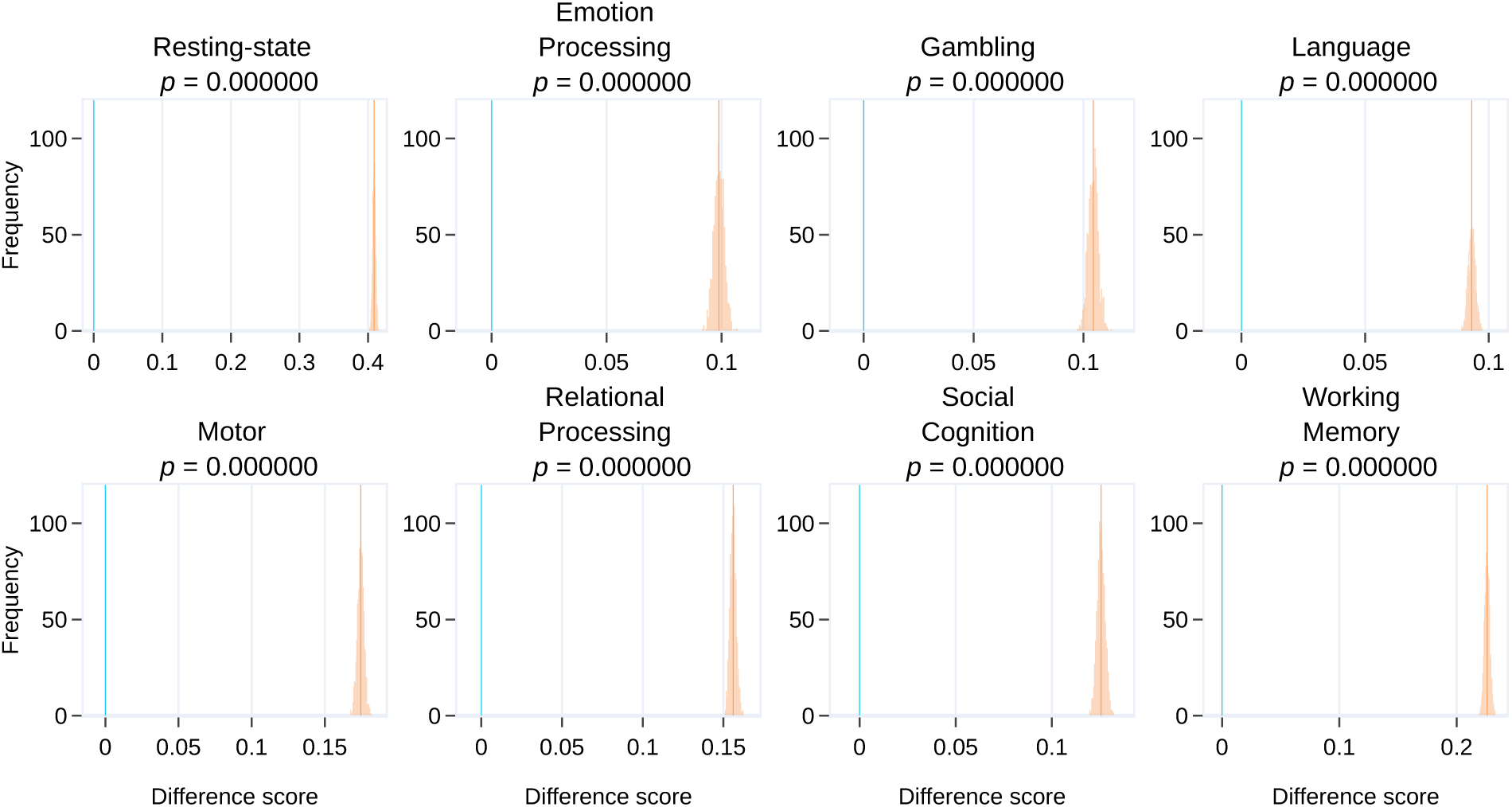
Combined subnetwork vs whole-cortex FCs: Comparison of identification accuracy based on combined subnetwork FCs (see Section 3.7) and whole-cortex FCs for each condition. The geodesic distance measure was used for identification. Data for each condition were trimmed such that they all had the same time course length (of 138; see Section 3.5). For each condition, the distributions shown in orange represent the difference between the mean participant identification accuracy based on combined subnetwork FCs and whole-cortex FCs across the outer bootstrap iterations (see Section 2.7). The orange line indicates the mean of the difference distribution and the blue line indicates zero difference.

## Footnotes

1 We thank one of the reviewers for helping solve this puzzle.

